# Targeted-SMN insufficiency in Skeletal Muscle Stem Cells mediates non-cell autonomous loss of motor neurons at long term

**DOI:** 10.1101/2024.12.19.629483

**Authors:** Jordan Mecca, Julien Mignot, Marianne Gervais, Teoman Ozturk, Stéphanie Astord, Juliette Berthier, Stéphanie Bauché, Julien Messéant, Maria-Grazia Biferi, Hélène Rouard, Martine Barkats, Frédéric Relaix, Nathalie Didier

## Abstract

Spinal Muscular Atrophy (SMA) is due to a deficit in SMN protein encoded by the *SMN1* gene. SMN-targeted disease modifying treatments have greatly improved the clinical outcomes of this neuromuscular disease. However, uncertainties remain regarding their long-term efficacy and non-neuronal tissue involvement in disease progression. We found that SMA type II patient muscles display a reduced number of quiescent PAX7+ Muscle Stem Cells (MuSC). In SMA mice, we showed that SMN is an important regulator of myogenic progenitor fate during early postnatal growth. In *Pax7 Cre*-driven conditional knockout mouse models, we demonstrated that high levels of SMN are required to ensure the maintenance of the quiescent MuSC pool in adult muscle. We further established that depletion of SMN-deficient MuSC yielded neuromuscular junctions remodeling followed by a non-cell autonomous loss of motor neurons in the long term. Overall, our findings demonstrate that MuSC are a crucial therapeutic target for SMA treatment.

**HIGHLIGHTS:**

- SMN regulates myogenic lineage progression and quiescent MuSC pool establishment during postnatal growth
- Both *Smn* alleles are necessary for the survival of quiescent MuSC in adult muscle
- Depletion of SMN-deficient MuSC leads to NMJ remodeling and non-cell autonomous loss of MN

## INTRODUCTION

Spinal Muscular Atrophy (SMA) is a devastating neuromuscular disorder that, although rare, is the leading genetic cause of infant mortality (incidence ∼ 1/10,000 live births) ^1^. This autosomal recessive disorder is characterized by the degeneration of motor neurons (MN) from the ventral horn of the spinal cord, muscle weakness and atrophy leading to premature death in the most severe forms. In most cases, SMA results from homozygous mutations or deletions in the *Survival of Motor Neuron 1* gene (*SMN1*) encoding SMN protein ^2^. In humans, a paralogous gene *SMN2* carrying a C>T transition in exon 7 is present. This transition leads to the exclusion of the exon 7 in 90% of *SMN2* transcripts and the production of an unstable truncated protein (SMNΔ7). In contrast, 10% of *SMN2* transcripts include exon 7 allowing the production of a functional full-length SMN protein ^3–5^. SMA is therefore caused by a deficit in functional SMN protein, which is exclusively produced by the *SMN2* gene. Accordingly, the number of *SMN2* copies is generally related to the severity of the disease ^6,7^. SMA is classified into different types (Type 0 to IV) according to the time of onset of the symptoms and the severity of the disease. Patients with SMA type I, the most frequent form of SMA, are usually carrying 2 copies of *SMN2*, develop symptoms before 6 months old and have a life expectancy of less than 2 years old without any treatment ^8^. In the last few years, major advances have been made in the treatment of SMA with the approval of three disease modifying treatments targeting SMN through different modes of action and routes of administration. Nusinersen, an antisense oligonucleotide delivered intrathecally, and risdiplam, an orally deliverable splicing modifier, both act by promoting the inclusion of exon 7 in *SMN2* transcripts during the splicing process ^9–12^. Onasemnogene abeparvovec, is an Adeno-Associated Viral vector of serotype 9 carrying *SMN1* gene delivered as a unique systemic injection, to replace the mutated *SMN1* gene ^13–15^. These 3 treatments have proven significant and unprecedented therapeutic benefits, in terms of survival and motor milestones, especially for children with a severe form of SMA treated early after birth ^10,12,15^. Despite these encouraging results, these therapies are not always curative and yield variable responses especially in patients with SMA types II, III and IV ^16^, highlighting the need to clarify the specific requirements for SMN and the long-term impacts of these therapies at the cellular and molecular levels, including in non-neuronal tissues. Indeed, it is well established that SMN protein levels are elevated during development and postnatal periods and then decline with age in humans and mice ^17,18^, consistent with the optimal therapeutic effect obtained with early treatment ^19^. However, whether there are specific requirements for SMN protein at the tissue and cellular level in non-neuronal tissues throughout life is less documented.

In addition, multiple lines of evidence indicate that therapeutic benefits require restoration of SMN expression in MN as well as in peripheral tissues. MN-targeted restoration of SMN ^20–24^ or inhibition of MN apoptosis ^25^ provided limited improvement in survival of SMA mice. Consistent with these findings, several peripheral tissues were shown to be affected by SMN deficiency, among which skeletal muscle tissue and its resident progenitor cells: Fibro-Adipogenic Progenitors and Muscle Stem Cells (MuSC) ^26–32^. SMN was found to interact with sarcomeric proteins of muscle fibers and disruptions of actin cytoskeleton associated with altered localization of SMN were reported in SMA type I muscles supporting an involvement of SMN in sarcomere architecture ^33–35^. In agreement, mutant mice carrying a deletion of *Smn* exon 7 restricted to muscle fibers exhibited a severe dystrophic phenotype and a reduced life expectancy ^36^. Moreover, the muscle phenotype of these mice was worsened when MuSC only had one copy of *Smn* ^28^. Early post-natal myogenesis defects were reported in SMA patient and SMA mouse muscles ^31,37–40^ whereas selective restoration of SMN under the control of *Myf5* and *MyoD* promoters in myogenic precursors of SMA mice improved their motor functions and survival ^21^. Lastly, SMN-deficient myoblasts exhibited impaired differentiation capacity *in vitro* ^41–44^. Altogether, these data support the hypothesis of a muscle-specific role for SMN, recently reinforced by a study demonstrating a primary role for SMN-deficient muscle tissue in the progression of SMA ^29^. However, a direct contribution of the MuSC, essential players in muscle homeostasis and regenerative potential, in the course of the disease remains to be established. In addition, we anticipate that an inefficient targeting of this tissue and cellular compartments by current treatments could, in the long-term, be detrimental to the neuromuscular system. In favor of this argument, a recent study revealed unresolved muscle defects in treated SMA type II patients that could underlie the variable response to treatment ^45^.

Here, we further investigated the role of SMN in the regulation of MuSC fate during early postnatal growth and in the adult muscle. We noted that the number of quiescent PAX7+ MuSC in muscles from juvenile SMA type II patients was significantly reduced compared to controls. We showed that SMN level regulates MuSC progression in the myogenic lineage and establishment of the quiescent MuSC pool during early postnatal growth. Using inducible conditional knockout mouse models, we observed that reduced levels of SMN rapidly induced quiescent MuSC apoptosis, leading to MuSC reservoir depletion in adult muscle. We showed that SMN-deficient MuSC depletion leads to Neuromuscular Junction (NMJ) remodeling within 2 weeks post-tamoxifen (TMX) injections. Moreover, we demonstrated that the depletion of SMN-deficient quiescent MuSC triggered a selective loss of part of the αMN in the spinal cord while γMN were not affected, 6 months post-TMX. Overall, our findings highlight a non-cell autonomous mechanism of MN loss that could participate in the natural course of intermediate (type II) and late-onset (type III and IV) SMA but also have important implications for the long-term treatment of patients.

## RESULTS

### Juvenile SMA type II muscle biopsies exhibit reduced number of quiescent PAX7+ MuSC

Early observations by electron microscopy reported a deregulation of MuSC number in SMA type I patient muscles compared to newborn control muscles ^37,38^. Later on, an increased number of PAX7+ MuSC was reported in newborn SMA type I patients supporting a defect in early myogenesis ^40^. In order to investigate the impact of SMN deficit in later stage disease, we quantified the number of PAX7+ MuSC in muscle biopsies obtained from SMA type II patients (SMA) and control donors (CTR) aged from 10 to 18 years (**Table 1**). We classified SMA biopsies according to the severity of their phenotype, *i.e.* heterogeneity in fiber size and extent of fibrosis and adipose tissue infiltration (**Fig 1A**). Our analyses revealed that the mean number of PAX7+ MuSC per field in each SMA patient muscle taken individually was reduced compared to the mean number of PAX7+ MuSC per field in CTR muscle group (**Fig 1B, C**). Similarly, the mean number of PAX7+ MuSC per field in SMA muscle group was significantly lower than in the CTR muscles (**Fig 1D**). By performing co-immunostaining for PAX7 and the proliferation marker KI67 (**Fig 1B**), we did not detect any PAX7+KI67+ cells in either SMA or CTR muscles, indicating that at this stage, PAX7+ MuSC are predominantly in a quiescent state in human muscles. Altogether, our observations suggest that the establishment and/or the maintenance of the quiescent MuSC pool is affected in SMA patient muscles. We thus sought to determine whether the reduced number of PAX7+ MuSC was due to an exhaustion of the pool as a consequence to the severe muscle phenotype and/or a direct loss of the MuSC linked to their intrinsic SMN-deficit.

**Figure 1:**
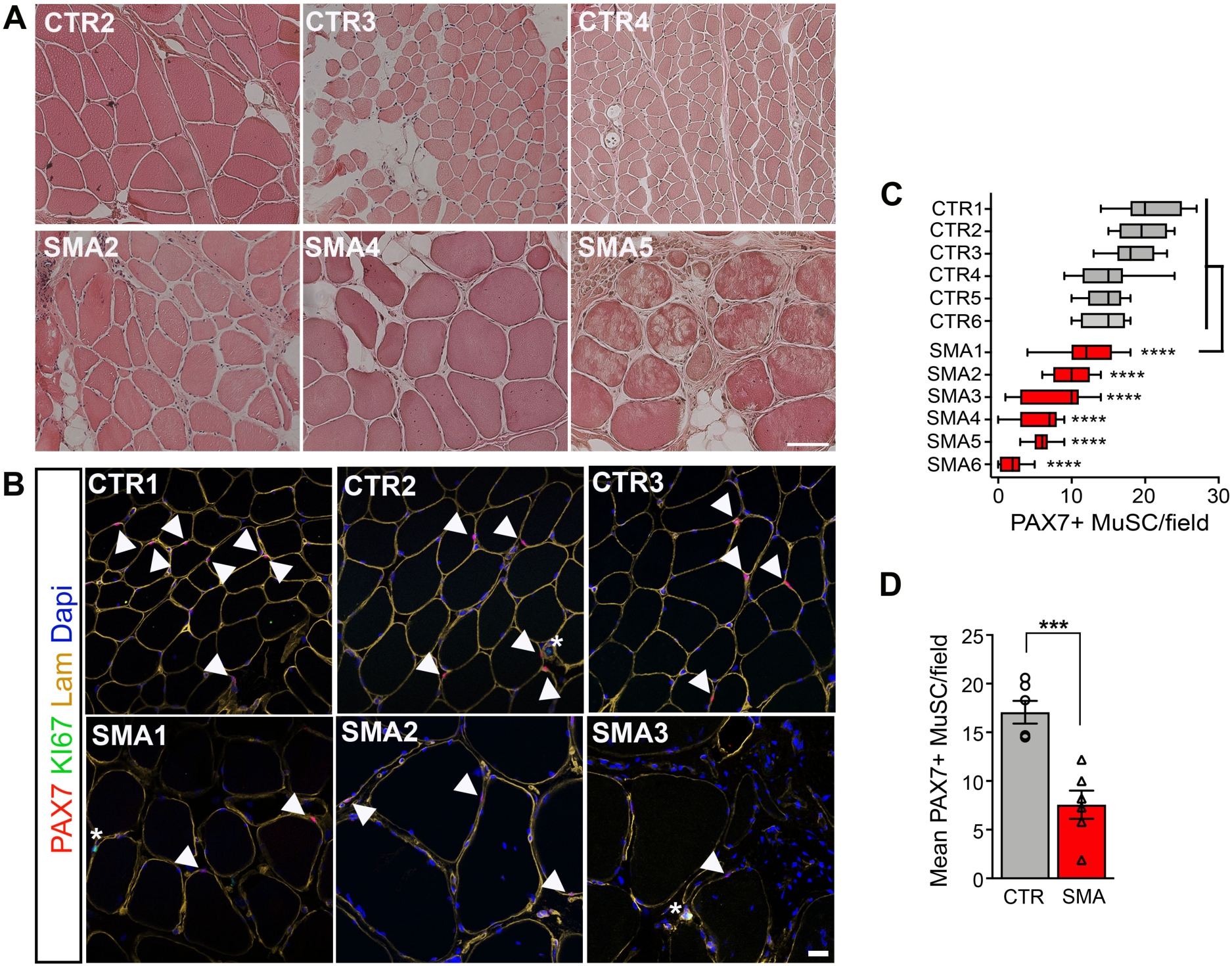
Muscles from juvenile SMA type II patients exhibit reduced PAX7+ MuSC pool. (**A, B**) Representative cross-sections of CTR and SMA muscles stained with hematoxylin and eosin (**A**), or immunostained for PAX7, KI67 and Laminin. Nuclei were stained with DAPI (**B**). Arrowheads point to PAX7+ cells. Stars indicate KI67+ cells. Scale bars: 500 μm and 20μm. (**C**) Box and whisker plots showing the distribution of PAX7+ cells per field. (**D**) Graph showing the mean number of PAX7+ cells/field. Data were normalized per field due to the high heterogeneity of fiber sizes, especially in SMA muscles. Values are the means±sem obtained from 6 different donors/group. One-way ANOVA and Dunnett’s multiple comparisons test (**C**), and Unpaired T test (**D**), with *\*\*\*p<0.001* and *\*\*\*\*p<0.0001*.

**Table 1:**
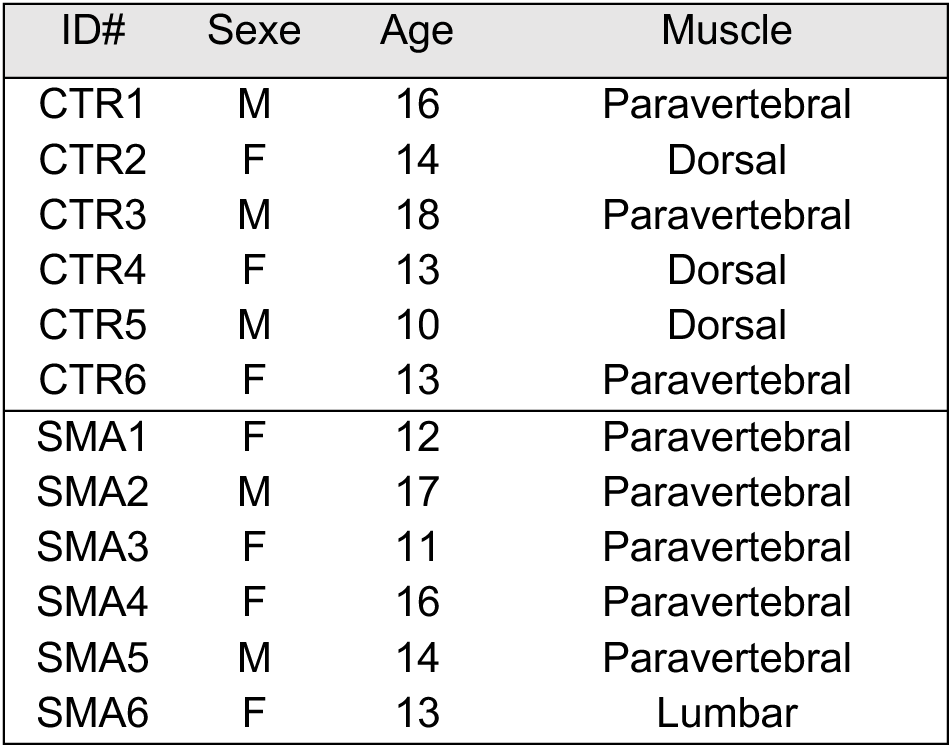
Characteristics of SMA type II patient muscle biopsies.

### SMN level regulates myogenic lineage progression of myogenic progenitors during early postnatal myogenesis

We first characterized the early postnatal myogenesis of the severe *hSMN2* SMA mouse model (**Fig 2**). In this model, previous study reported a dysregulation of myogenic cell behavior in new born mice (P1) ^31^. To further investigate the role of SMN in the myogenic lineage progression, we examined the expression of the myogenic factors PAX7, MYOD and Myogenin (MYOG) in *Smn^-/-^;hSMN2* (SMA) and *Smn^+/+^;hSMN2* (CTR) mouse muscles through immunofluorescence at the P4 stage of the postnatal myogenesis (**Fig 2A-C**). We quantified the number of cells PAX7+MYOD- (quiescent MuSC), PAX7+MYOD+ (activated progenitors), PAX7-MYOD+ (committed precursors) and MYOG+ (differentiating myocytes). We observed a significant increase of the total number of PAX7+ cells per 100 fibers in SMA mouse muscles relative to controls, which mainly resulted from an increased number of PAX7+MYOD+ progenitors (**Fig 2B**). Conversely, the number of PAX7+MYOD- quiescent MuSC (**Fig 2B**) or differentiating MYOG+ cells (**Fig 2C**) was reduced in SMA mouse muscles relative to controls. These observations revealed that SMN-deficit affects the fate of myogenic cells during early postnatal myogenesis. In particular, our data suggested that insufficient levels of SMN keep myogenic cells in an activated state affecting their ability to downregulate MYOD to engage in quiescence and/or to downregulate PAX7 to progress in the differentiation program ^46^.

**Figure 2:**
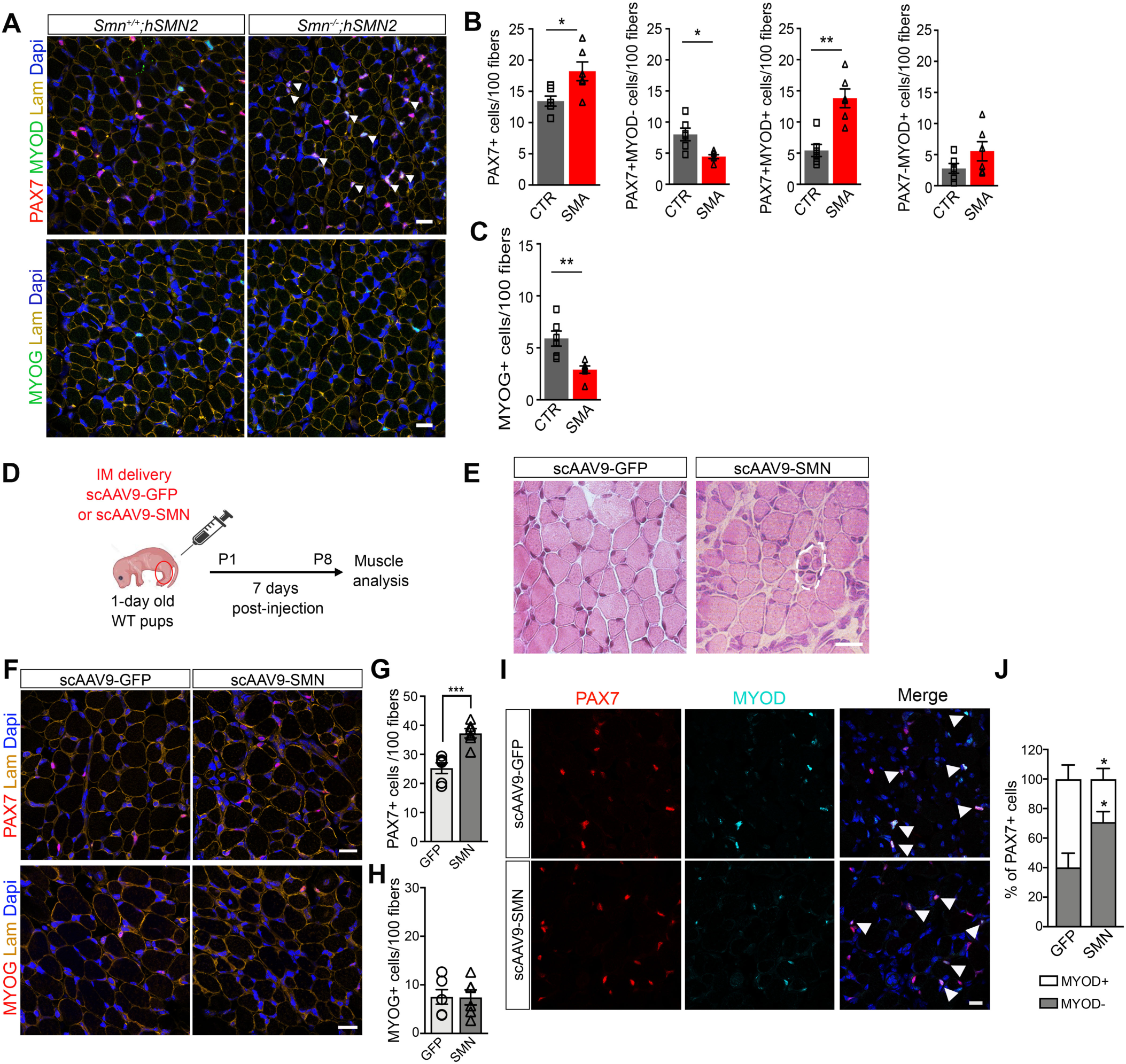
SMN regulates myogenic progenitor progression in the myogenic lineage program during early postnatal growth. (**A**) Representative cross-sections of Tibialis Anterior (TA) muscles from 4-days old *Smn^+/+^;hSMN2* (CTR) and *Smn^-/-^ ;hSMN2* (SMA) mice immunostained for PAX7, MYOD and Laminin or MYOG and Laminin. White arrowheads indicate PAX7+MYOD+ cells. Nuclei were stained with DAPI. Scale bars: 20μm. (**B, C**) Quantification of the number of total PAX7+, PAX7+MYOD-, PAX7+MYOD+, PAX7-MYOD+ cells (**B**), and the number of MYOG+ cells (**C**) normalized per 100 fibers. Values are means±sem of 6 independant quantifications (n=6 mice/group). Unpaired T test with *\*p<0.05* and *\*\*p<0.01*. (**D**) Experimental chronograph. IM: Intramuscular. (**E**) Representative cross-sections of TA muscles stained with Hematoxylin/Eosin at P8. Dotted lines show a cluster of small fibers with centro-located nuclei. (**F**) Representative cross-sections of TA muscles immunostained for PAX7 or MYOG and Laminin at P8. (**G, H**) Quantification of the number of PAX7+ cells (**G**), or MYOG+ cells (**H**) per 100 fibers. Unpaired T test with *\*\*\*p<0.001*. (**I**) Representative cross-sections of TA muscles immunostained for PAX7 and MYOD. Arrowheads point to PAX7+MYOD+ cells. Nuclei were stained with DAPI. Scale bars: 20μm. (**J**) Quantification of the proportion of PAX7+ cells MYOD- (grey bars) or MYOD+ (white bars). Values are means±sem of 6 independent quantifications (n=6 mice/group). Two-way ANOVA and Sidak’s multiple comparison test with *\*p<0.05*.

To further explore the role of SMN in the myogenic lineage progression, we next overexpressed SMN in 1 day old wild-type mouse muscles, using intramuscular delivery of a scAAV9-SMN vector (self-complementary Adeno Associated Virus of Serotype 9 carrying the human *SMN1* cDNA under the control of the ubiquitous PGK promoter) or a scAAV9-GFP vector as a control (**Fig 2D**). In our experimental conditions, a mean of 90% of the MuSC were transduced (**Fig S1A, B**). Furthermore, using immunostaining on sections from injected muscles, we could confirm the overexpression of SMN in PAX7+ cells (**Fig S1C**). Macroscopic analysis revealed the presence of clusters of small myofibers with centro-located nuclei in scAAV9-SMN injected muscles, showing that SMN overexpression disrupted the postnatal growth process (**Fig 2E**). We next analyzed PAX7 and MYOG expression by immunofluorescence, 7 days after delivery (**Fig 2F**). We observed a significant increase of the total number of PAX7+ cells/100 fibers in muscles injected with scAAV9-SMN vector compared to muscles injected with scAAV9-GFP control vector (**Fig 2G**), whilst the number of differentiating MYOG+ cells was similar (**Fig 2H**). Furthermore, the proportion of PAX7+ cells co-expressing MYOD (PAX7+MYOD+) was significantly reduced in muscles injected with scAAV9-SMN vector relative to controls (**Fig 2I, J**). In contrast to SMN deficiency, these observations suggested that an excess of SMN during early postnatal growth may instead favor the commitment of myogenic progenitors to quiescence at the expense of differentiation. Altogether, our data demonstrated that SMN levels have to be finely regulated to ensure proper progression of myogenic cells through the myogenic program and the establishment of the quiescent MuSC pool during postnatal myogenesis.

### MuSC-targeted deletion of a single or both *Smn* alleles induces rapid apoptosis leading to quiescent MuSC pool depletion in adult muscle

SMN is involved in multiple essential cellular functions, making it necessary for cell survival ^47^. In addition, the presence of apoptotic MuSC has been reported in SMA mouse muscles suggesting that reduced level of SMN could affect MuSC viability ^48^. To further understand the mechanisms underlying the reduction in the number of quiescent MuSC in SMA patient muscles, we next sought to investigate the role of SMN in quiescent MuSC pool maintenance. To this end, we made use of an inducible conditional knockout (cKO) mouse model *Pax7^CreERT2/+^;Smn* floxed in which *Smn* floxed allele deletion is driven by the inducible CreERT2 recombinase under the control of the *Pax7* regulatory regions ^49,50^. We analyzed apoptosis at early time points following TMX injections by flow cytometry (**Fig 3A**). In brief, mononucleated cells from adult *Pax7^CreERT2/+^;Smn^+/+^* (CTR), *Pax7^CreERT2/+^;Smn^fl/+^* (HTZ) and *Pax7^CreERT2/+^;Smn^fl/fl^*(HMZ) digested muscles were immunostained for specific surface markers to select the MuSC population (Lin^-^SCA1^-^CD34^+^ITGA7^+^) and stained with coupled-Annexin V to detect apoptotic cells (**Fig 3B-D**). These analyses revealed that after 4 injections of TMX (T3), HTZ and HMZ muscles exhibited an increased proportion of early apoptotic MuSC (defined as AnnexinV^+^7AAD^-^ cells) compared to CTR muscles (**Fig 3E**). In contrast, the proportion of early apoptotic Lin^-^SCA1^+^ cells was stable and similar for the 3 genotypes between T0 and T3, confirming that this increased apoptosis was specifically mediated by the Cre-driven recombination in PAX7+ MuSC (**Fig S2A, B**). We next quantified the number of PAX7+ MuSC on muscle sections, 1-month post-TMX (**Fig 3F, G**). CTR, HTZ and HMZ muscles displayed the same number of PAX7+ MuSC/100 fibers prior to TMX injections (**Fig S2C**). In contrast, we observed a marked depletion of the PAX7+ MuSC pool in HTZ and HMZ muscles (60% and 76%, respectively) as compared to CTR muscles following TMX injections (**Fig 3G**). Importantly, we generated another cKO model (referred as *Pax7^CreERT2(Gaka)^;Smn* floxed mice), in which the CreERT2 sequence is inserted downstream of the *Pax7* stop codon allowing normal endogenous PAX7 expression ^51^. In this model, we observed comparable rates of MuSC depletion (60% and 71% in HTZ and HMZ, respectively, **Fig S2D**), demonstrating that MuSC loss was not due to *Pax7* haploinsufficiency ^52^. Our findings demonstrate that high levels of SMN are necessary for the survival of PAX7+ MuSC and thereby for the maintenance of quiescent MuSC reservoir in adult muscle. We assume that the different level of MuSC depletion between HTZ and HMZ muscles in both cKO mouse models could result from different recombination efficiencies, or from a heterogeneity in SMN protein requirements among the MuSC pool. Supporting this hypothesis, available single cell RNA sequencing (scRNAseq) data show that murine and human MuSC display heterogeneous levels of expression of *Smn* gene (**Fig S2E-J**). Importantly, we noticed that part of HMZ and in a lesser extent of HTZ mice, developed a severe phenotype (weight loss, kyphosis and limb paralysis) and ultimately died between 14 and 30 weeks post-TMX injections suggesting neuromuscular system damage (**Fig 3H**).

**Figure 3:**
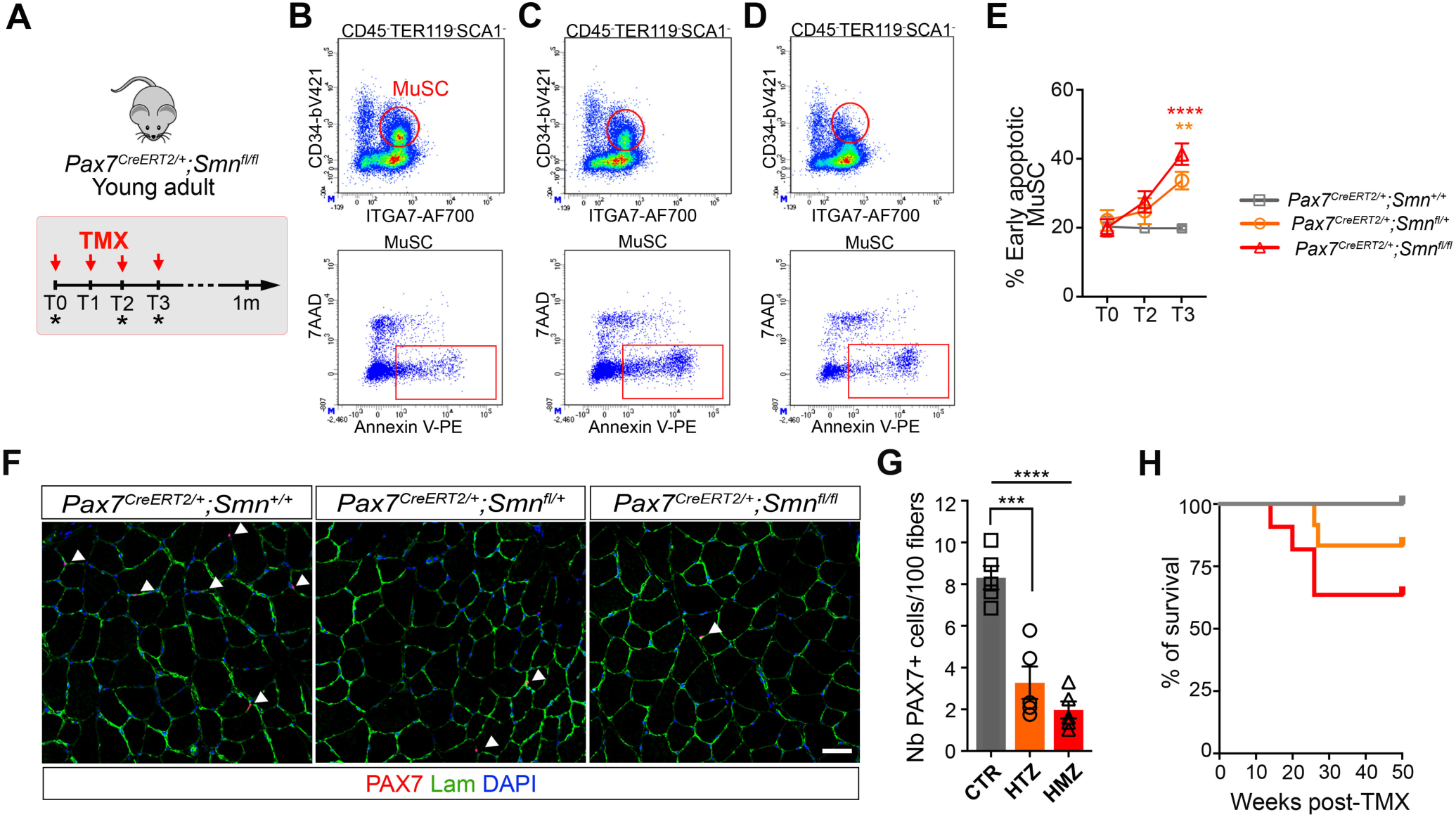
Loss of a single or both *Smn* alleles leads to rapid apoptosis and depletion of quiescent MuSC pool in adult muscle. (**A**) Experiment chronograph. (**B-D**) Representative scatter plots showing the strategy used to quantify the proportion of apoptotic MuSC (identified as CD45-TER119-SCA1-CD34+ITGA7+ cells) among mononucleated cells of *Pax7^CreERT2/+^;Smn^+/+^* (CTR, **B**), *Pax7^CreERT2/+^;Smn^fl/+^*(HTZ, **C**) and *Pax7^CreERT2/+^;Smn^fl/fl^* (HMZ, **D**) digested muscles, after 4 injections of TMX (T3). (**E**) Graphs showing the proportion of early apoptotic (AnnexinV+7AAD-) MuSC. Data are the mean±sem of 3 to 4 independent experiments. Two-way ANOVA and Tukey’s multiple comparisons test with *\*\*p<0.01* and *\*\*\*\*p<0.0001*. (**F**) Representative TA cross-sections from adult CTR, HTZ and HMZ mice immunostained for PAX7 and Laminin, 1-month post-TMX injections. Nuclei were stained with DAPI. Arrowheads indicate PAX7+ MuSC. Scale bar: 40 μm. (**G**) Quantification of the number of PAX7+ MuSC per 100 fibers. Data are mean±sem of 5 independent quantifications (n=5 mice/group). One-way ANOVA and Tukey’s multiple comparisons test with *\*\*\*p<0.001* and *\*\*\*\*p<0.0001*. (**H**) Kaplan Meier survival curve (10-12 mice/genotype). Log-rank (Mantel-Cox) test (CTR *vs* HMZ, *p=0.08*).

### *Pax7^CreERT2/+^;Smn^fl/+^* and *Pax7^CreERT2/+^;Smn^fl/fl^*mouse muscles exhibit signs of denervation/reinnervation

Neuromuscular disruptions are usually associated with myofiber atrophy and abnormal profiles of MyHC isoform expression ^53^. We therefore measured fiber cross-sectional areas (CSA) and carried out a phenotypic characterization of TA muscles, 1 and 6 months post-TMX (**Fig 4A, B**). One-month post-TMX, we noticed a reduction in fiber size of HMZ mouse muscles compared to HTZ and CTR muscles whereas fiber composition was unchanged, consistent with a denervation event (**Fig 4C, D**). At 6 months post-TMX injections, the myofiber CSA were equivalent in the 3 genotypes (**Fig 4E**). However, we observed a significant increase in the proportion of type IIa fibers associated with a reduction in type IIb fibers in HMZ relative to CTR muscles, a typical feature of a reinnervation process. (**Fig 4F**). Of note, a similar tendency of fiber phenotype changes was observed in HTZ muscles presumably due to a slower kinetic of denervation-reinnervation than in HMZ (**Fig 4F**).

**Figure 4:**
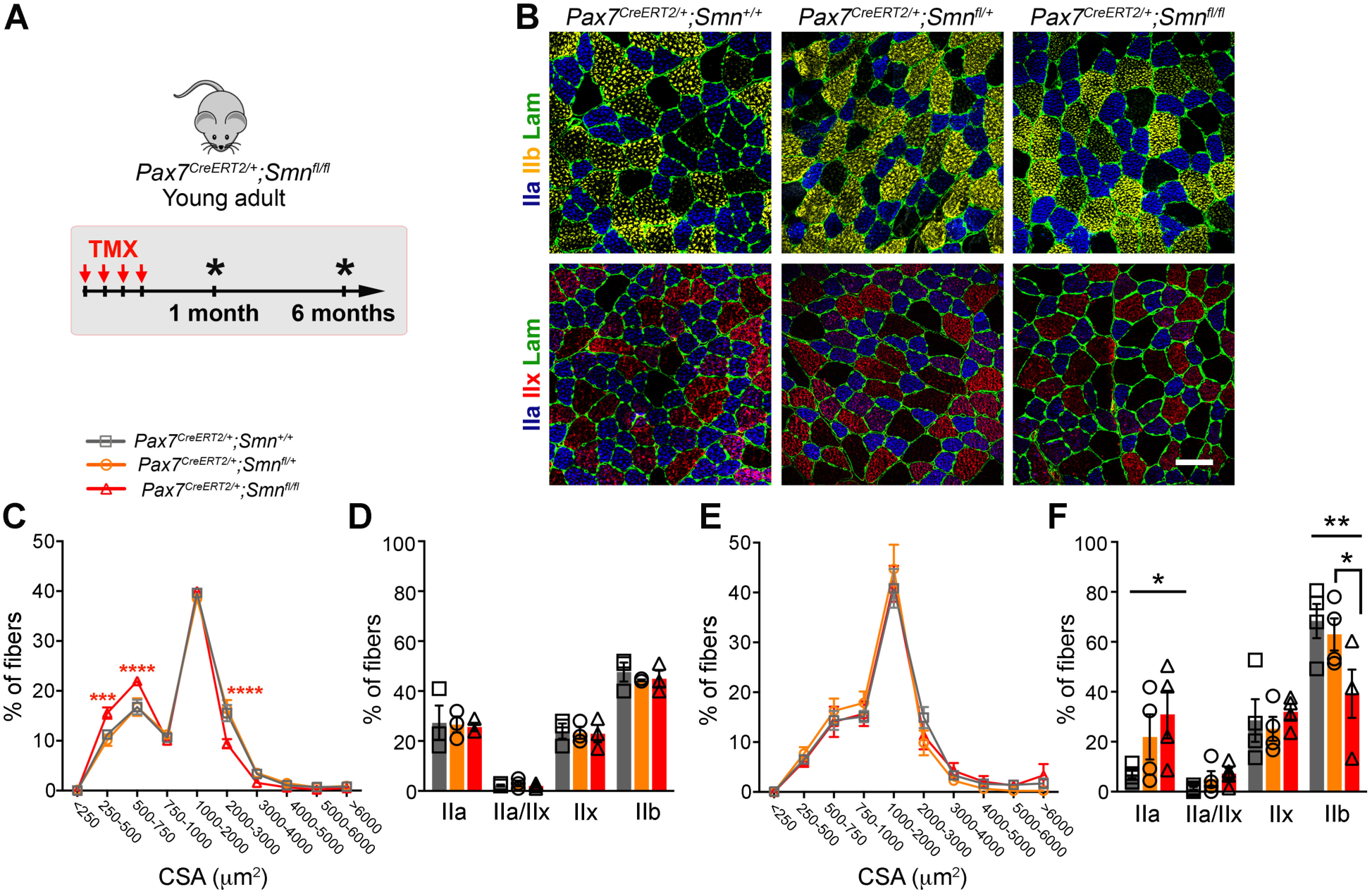
Targeted depletion of SMN-deficient MuSC triggers phenotypic switch of muscle fibers. (**A**) Experiment chronograph. (**B**) Representative cross-sections of TA muscles from CTR, HTZ and HMZ mice, immunostained for MyHC IIa and IIb or IIx and Laminin, 1-month post-TMX injections. Scale bar:100μm. (**C-F**) Graphs showing the distribution of fiber Cross-Sectional Areas (CSA) and the proportion of each fiber types, 1-month post-TMX (**C**, **D**) and 6 months post-TMX (**E**, **F**). Values are the means±sem of 3 to 4 independent quantifications (n=3 mice/genotype for 1-month post-TMX, n=4 mice/genotype for 6 months post-TMX data). Two-way ANOVA and Tukey’s multiple comparisons test with *\*p<0.05* and *\*\*p<0.01*.

### Depletion of SMN-deficient MuSC leads to NMJ remodeling and non-cell autonomous loss of MN throughout the spinal cord at long term

SMA is characterized by the selective degeneration of the motor neurons (MN) of the ventral horn of the spinal cord. Muscle fibers and MN are closely connected through the neuromuscular junctions (NMJ). Previous studies using murine models of MuSC depletion have established a role of MuSC in the proper regeneration and maintenance of the NMJ upon neuromuscular disruptions and aging ^54,55^, and a sub-population of NMJ-associated MuSC responsive to denervation was identified ^56^. Moreover, recent study reported a reduction of the number of subsynaptic myonuclei in *Smn^2B/-^* mouse model presumably due to a decrease number of MuSC ^32^.

By 3D imaging on cleared TA muscles from *Pax7-nGFP* mice, we observed that the density of MuSC (number of MuSC/mm^3^) was significantly higher in the vicinity of the NMJ (**Fig 5A, B**) and that 35% of the MuSC are located at less than 25μm from a NMJ (**Fig 5C**), in agreement with previous study ^54^. Given the physical and functional link between the MuSC and the NMJ, we sought to investigate whether SMN-deficient MuSC depletion could have an impact on the MN through the NMJ. We first verified the absence of leaky Cre activity in non-injected mice and of unanticipated Cre activity in MN or in neighboring cells using a reporter mouse model *Pax7^CreERT2/+^;R26^mTmG^*^57^ (**Fig S3**). In these reporter mice, all the cell membranes constitutively express the *Tomato* transgene, whereas the *GFP* transgene expression is restricted to the cells in which a Cre-mediated recombination event occurred. No GFP expression was observed in muscles and spinal cords of non-injected mice, confirming the absence of leaky activity of the Cre recombinase in our model. We then injected TMX for 4 consecutive days to adult *Pax7^CreERT2/+^;R26^mTmG^*and *Pax7^+/+^;R26^mTmG^* mice and examined GFP expression by immunofluorescence on spinal cord and TA muscle sections, 5 days later (**Fig S3A**). We detected robust GFP expression on TA muscle sections resulting from Cre-driven recombination in PAX7+ MuSC (**Fig S3B**). Conversely, no GFP expression was detected at any level of the spinal cord (*i.e.* cervical, thoracic and lumbar levels) confirming that no recombination event occurred in MN, nor in neighboring cells (**Fig S3C**). Moreover, we took advantage of available single nuclei RNAseq data performed on adult mouse spinal cord to confirm the absence of expression of *Pax7* gene in MN (**Fig S3D, E**).

**Figure 5:**
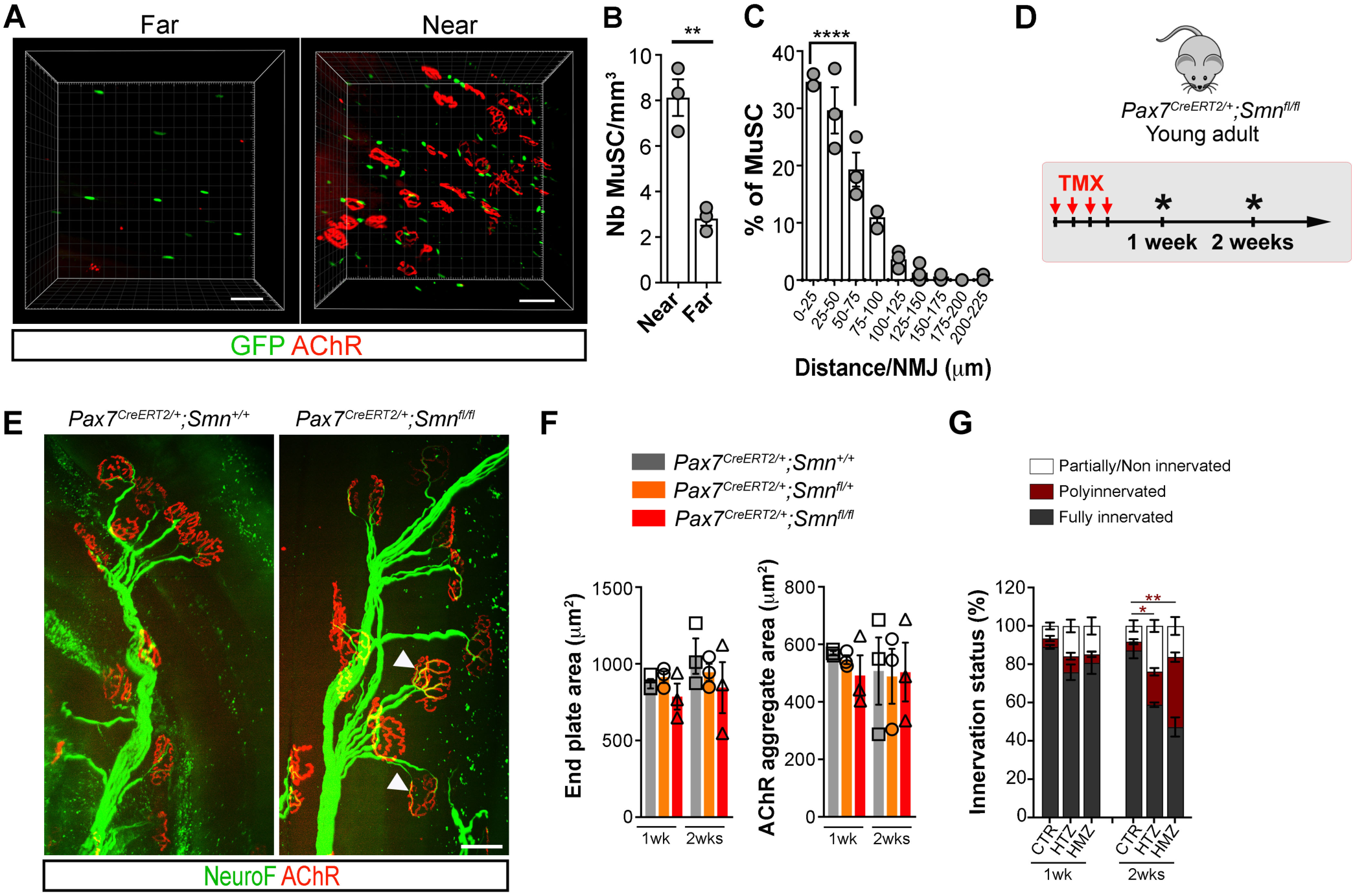
Depletion of SMN-deficient MuSC induces NMJ remodeling. (**A**) 3D imaging of NMJ and MuSC on cleared TA muscle of adult *Pax7-nGFP* transgenic mouse. AChR clusters were labeled with coupled α-Bungarotoxin (α-BTX). Scale bars: 50 μm. (**B, C**) Graphs showing the density of MuSC located near (≤100μm) or far (>100 μm) from a NMJ (**B**), the distance between MuSC and NMJ on a muscle fiber (**C**). Values are the mean±sem from 3 independent experiments. Unpaired T test with *\*\*p<0.01* and one-way ANOVA and Dunnett’s multiple comparisons test with *\*\*\*\*p<0.0001*. (**D**) Chronograph of the experiment. (**E**) Representative images of maximum z-projections showing NMJ in TA muscles 2 weeks post-TMX injections. Post-synaptic compartment was labeled with coupled α-BTX, pre-synaptic compartment was immunostained for neurofilaments (NeuroF). Scale bar: 50 μm. Arrowheads point to polyinnervated NMJ. (**F**) Morphological analysis of the NMJ. (**G**) Innervation status of the NMJ. Values are the mean±sem of 3 independent quantifications (n=3 mice/genotype). Two-way ANOVA and Tukey’s multiple comparisons test with *\*p<0.05* and *\*\*p<0.01*.

Having confirmed that Cre-driven recombination was restricted to MuSC in our model, we then examined the morphology and the innervation status of the NMJ in CTR, HTZ and HMZ mouse muscles, 1 and 2 weeks post-TMX injections (**Fig 5D-G**). We did not notice significant morphological changes of the NMJ between the 3 genotypes, at 1 and 2 weeks post-TMX (**Fig 5F**). However, we observed a slight increase in the proportion of partially/denervated NMJ at 1 week, which became more pronounced at 2 weeks in HTZ muscles compared with CTR. In addition, the proportion of poly-innervated NMJ was significantly increased in HTZ and HMZ muscles relative to CTR at 2 weeks, demonstrating that SMN-deficient MuSC ablation rapidly induced NMJ pre-synaptic compartment remodeling or instability (**Fig 5G**).

We next examined the impact of this NMJ remodeling on MN. For that, we characterized the MN pool in the ventral horn of the spinal cords of CTR, HTZ and HMZ mice, 1 and 6 months post-TMX (**Fig 6A-C**). No difference between the 3 genotypes was observed in non-injected mice (**Fig S4A**). One-month post-TMX injections, SMN expression was detectable in all the ChAT+ MN (**Fig 6B**), and the number of MN was equivalent between the 3 genotypes (**Fig 6D**). Of note, we noticed a small decrease in MN in the HMZ cervical and lumbar spines, which was more pronounced in *Pax7^CreERT2(Gaka)^;Smn^fl/fl^* mice (**Fig S4B**). At 6 months post-TMX, our analyses revealed a reduction of 20-30% of the number of MN in HTZ and HMZ mice compared to CTR at all the levels of the spinal cord and in both cKO models (**Fig 6E** and **Fig S4C**). In addition, we assessed the motricity performance by rotarod test between 4 and 18-weeks post-TMX (**Fig S4D, E**) and observed similar latency to fall between the 3 genotypes, demonstrating that upper levels of the central nervous system (*i.e.* motor cortex and the cerebellum) were not affected. Lastly, to exclude a potential indirect effect due to *Smn* deletion in cell from other tissues expressing *Pax7*, we generated *Pax7^CreERT2(Gaka)^;R26^iDTR/iDTR^* mice enabling localized ablation of MuSC by Diphteria Toxin (DT) intramuscular injection (**Fig 6F-H**). DT was injected in Tibialis Anterior, Quadriceps and Gastrocnemius muscles the day after TMX injections and 3 months later (**Fig 6F**), leading to a strong reduction of the number of PAX7+ MuSC in *Pax7^CreERT2(Gaka)^;R26^iDTR/iDTR^*muscles compared to controls (**Fig S5A-C**). After 6 months, the number of MN in the cervical and the thoracic spinal cords was not affected (**Fig 6H**). In contrast, the number of MN in the lumbar spine, which innervate hindlimb muscles, was reduced in *Pax7^CreERT2(Gaka)^;R26^iDTR/iDTR^*mice compared to control mice (**Fig 6H**). Overall, our findings reveal an unanticipated interdependence between quiescent MuSC and MN reservoirs. Indeed, we demonstrated that damage to the quiescent MuSC reservoir in adult muscle has a direct impact on part of the spinal cord MN in the long term and mediates a non-cell autonomous loss of these cells.

**Figure 6:**
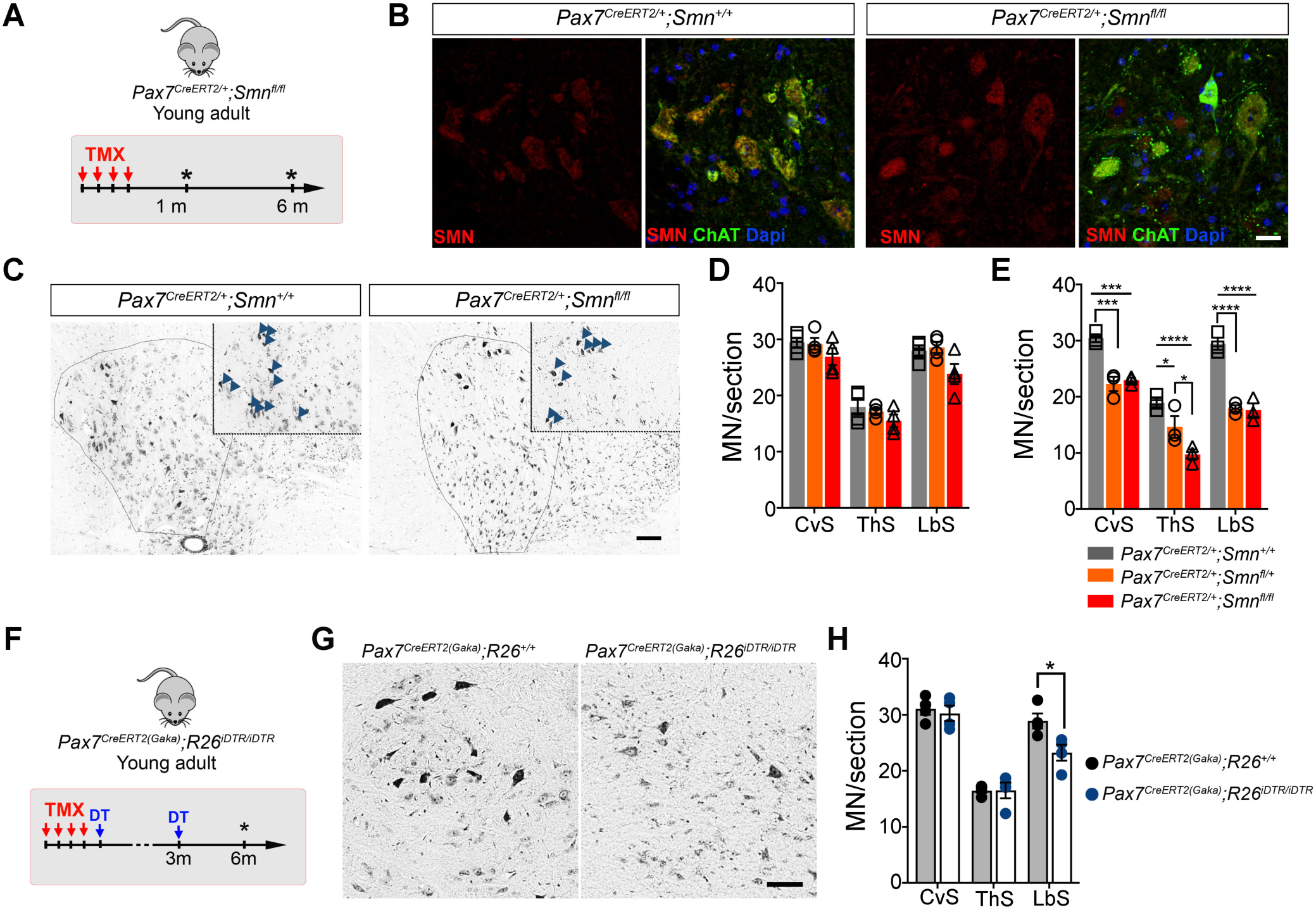
Depletion of SMN-deficient MuSC induces a non-cell autonomous loss of MN at long term. (**A**) Experiment chronograph. (**B, C**) Representative cross-sections of cervical spinal cords from CTR and HMZ mice immunostained for SMN and ChAT 1-month post-TMX (**B**) or stained with Niels coloration 6 months post-TMX (**C**). Scale bars: 25 and 100 μm, respectively. (**D, E**) Quantification of the number of MN/section,1-month (**D**) and 6 months post-TMX (**E**). CvS: Cervical Spine, ThS: Thoracic Spine and LbS: Lumbar Spine. Values are the means±sem of minimum 3 independent quantifications (3-4 mice/genotype). Two-way ANOVA and Tukey’s multiple comparisons test with *\*p<0.05*, *\*\*\*p<0.001* and *\*\*\*\*p<0.0001*. (**F**) Experiment chronograph. Diphteria Toxin (DT) was injected intramuscularly in Tibialis anterior, Gastrocnemius and Quadriceps muscles of adult *Pax7^CreERT2(Gaka)^;R26^+/+^*and *Pax7^CreERT2(Gaka)^;R26^iDTR/iDTR^* mice. (**G**) Representative images of lumbar spine cross-sections stained with Niels coloration, 6 months post-TMX. Scale bar: 50 μm. (**H**) Quantification of the number of MN/section. Values are represented as the means±sem of 4 independent experiments (n=4 mice/group). Two-way ANOVA and Sidak’s multiple comparisons test with *\*p=0.012*.

### Depletion of SMN-deficient MuSC selectively affects αMN

Our observation that SMN deficiency in quiescent MuSC may have long-term effects on MN integrity prompted us to further examine the profile of the affected MN. Within the mammalian spinal cord, lower MN pools in the ventral grey horn contain both alpha MN (aMN), which innervate extrafusal muscle fibers responsible for skeletal muscle contraction, and gamma MN (yMN), which innervate the intrafusal fibers of muscle spindles and regulate their sensitivity to stretch ^58^. aMN can be identified by the expression of both Choline AcetylTransferase (ChAT) and Neuronal Nuclei (NeuN), whereas yMN lack the expression of NeuN ^59,60^. We thus carried out immunohistochemical analysis on spinal cord sections of CTR, HTZ and HMZ mice labelled for ChAT and NeuN, 6 months post-TMX injections (**Fig 7A, B**). We quantified the number of aMN (ChAT+NeuN+) and yMN (ChAT+NeuN-) per section at the different levels of the spinal cord. We noticed a significant decrease in the number of aMN in HTZ and HMZ mouse spinal cord relative to CTR at the cervical, thoracic and lumbar levels (**Fig 7C**) whereas the number of yMN was unaffected (**Fig 7D**). Thus, we demonstrate that SMN deficit in quiescent PAX7+ MuSC can selectively impact aMN without affecting yMN as previously reported in SMA mice ^61^.

**Figure 7:**
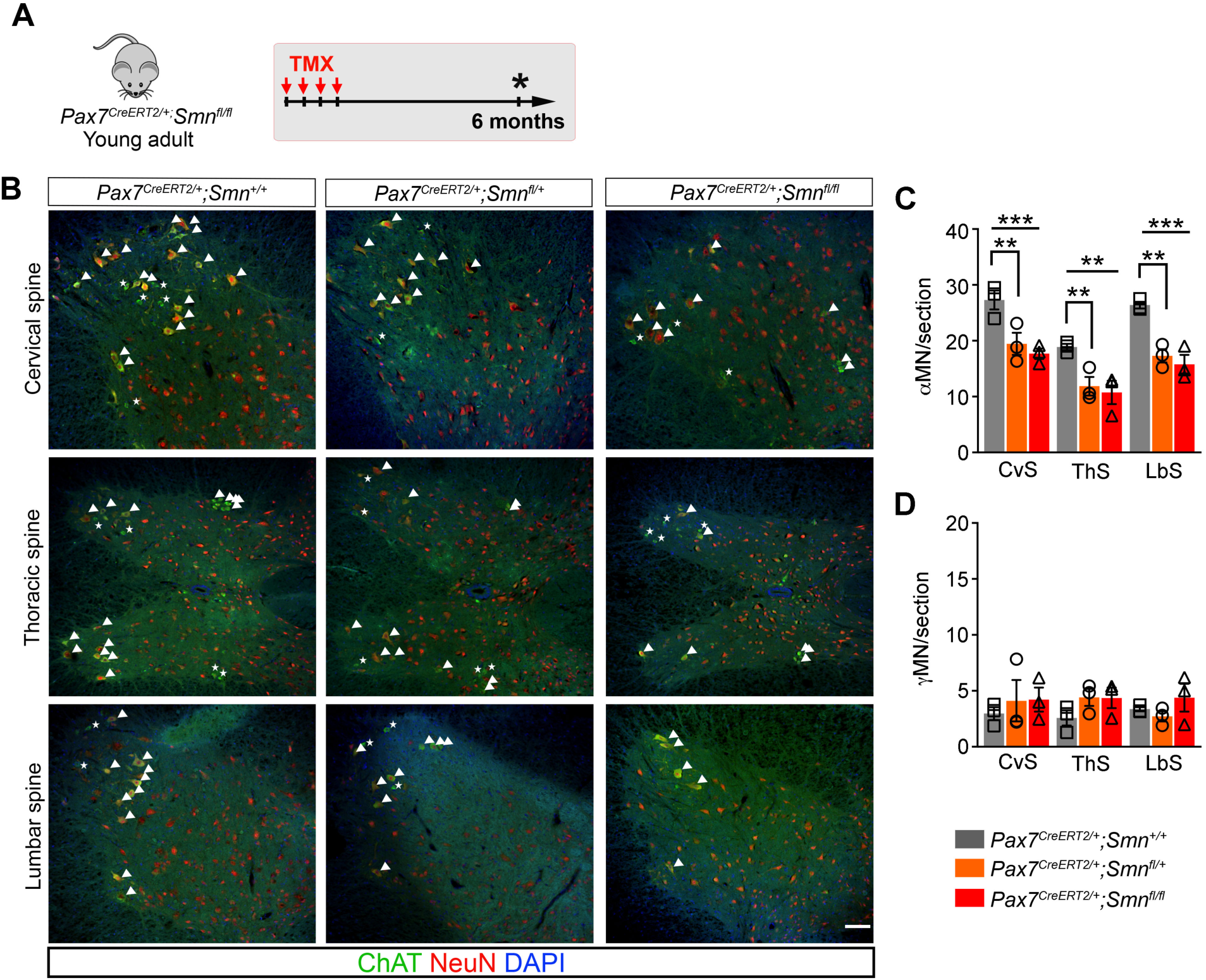
Depletion of SMN-deficient quiescent MuSC induces selective loss of αMN. (**A**) Experiment chronograph. (**B**) Representative cross-sections of spinal cords from Pax7*CreERT2/+_;Smn_+/+*_, *Pax7*_*CreERT2/+_;Smn_fl/+* _and *Pax7*_*CreERT2/+_;Smn_fl/fl* _mice_ immunostained for ChAT and NeuN, 6 months post-TMX. Nuclei were stained with DAPI. Arrowheads show αMN, stars show γMN. Scale bar: 100 μm. (**C, D**) Bar graphs showing the quantification of αMN (ChAT+NeuN+) and γMN (ChAT+NeuN-) per section. Values are the means±sem of 3 independent quantifications (3 mice/genotype). Two-way ANOVA and Tukey’s multiple comparisons test with *\*\*p<0.01* and *\*\*\*p<0.001*.

## DISCUSSION

It is well established that MN are the primary cellular targets of SMA. Nevertheless, growing evidence points to the importance of restoring SMN expression in peripheral tissues including muscle tissue to achieve satisfactory motor function and life span recovery ^23–25,45,62^. At the time of the first therapeutic successes with gene therapy ^10,15,19^, our findings reveal that MuSC are important therapeutic targets for the long-term preservation of the neuromuscular system, introducing the pathophysiological concept of SMA as a satellite cell-opathy.

Previous evidence suggested that SMN deficit could affect muscle tissue and MuSC behavior ^21,28,31,43,63^. However, a primary role of skeletal muscle in the progression of pathology has long been controversial ^20,21,29,64^ until it was clearly demonstrated that persistent SMN deficit specifically targeted to skeletal muscle tissue leads to the development of late-onset myopathy with NMJ dysfunctions and reduced motor performance ^29^. Moreover, transcriptomic analysis of muscle from SMA type II patients treated with nusinersen and/or risdiplam unveiled unaddressed mitochondrial abnormalities that sustain the inter-individual variable response to these treatments ^45^. Since the quiescent MuSC reservoir is necessary to ensure muscle tissue homeostasis and repair potential ^65,66^, it has thus become crucial to clarify the specific role and requirement for SMN in these cells to design next generation therapeutic strategies.

Post-mortem analyses of neonatal muscle biopsies of SMA type I patients reported an increased number of MuSC suggesting a defect in perinatal myogenesis ^37,38,40^. Our data reveal that at a later stage of the postnatal growth, the pool of quiescent MuSC is reduced in SMA patient muscles. We reasoned that this attrition of the quiescent MuSC pool could be a consequence of the severe phenotype of SMA muscles or a direct result of SMN deficit in MuSC. By characterizing early postnatal myogenesis of the severe SMA mouse model *hSMN2*, we observed an increased number of activated progenitors associated with a reduced number of PAX7+MYOD- quiescent MuSC and of differentiating cells. These observations are in agreement with previous studies showing that SMN-deficient myogenic cells display altered differentiation capacities ^31,41,43,44,63,67,68^. In addition, AAV-mediated SMN excess during early postnatal growth promoted the establishment of the quiescent MuSC pool at the expense of their differentiation. We thus assume that the previously reported increased number of MuSC in neonate SMA muscles could result from an increase in their proliferation rate driven by SMN deficit as shown for neural stem cells from SMA mouse embryos and SMA patient-derived iPSC ^69,70^. Conversely, at later stages of postnatal growth, the reduction in quiescent MuSC that we showed in SMA patient muscles may be partly elicited by an altered ability of SMN-deficient progenitors to enter into quiescence. Overall, our data highlight SMN as an important regulator of MuSC fate during postnatal growth, especially regarding their ability to enter into quiescence or to differentiate. Using conditional inducible KO mouse models, we evaluated the impact of SMN deficit in the maintenance of the adult quiescent MuSC pool. Consistent with the well-established anti-apoptotic role of SMN ^25,71–73^, the complete removal of SMN resulted in a rapid and robust induction of apoptosis in MuSC. Remarkably, ablation of a single *Smn* allele was sufficient to induce apoptosis and subsequent depletion of the PAX7+MuSC reservoir. This finding demonstrates that in adult muscle, high levels of SMN are essential to maintain of the quiescent MuSC pool and consequently the muscle’s regenerative potential.

Skeletal muscle myofibers and MN are closely linked via the NMJ, where MuSC are present at high density. A recent study identified a subset of MuSC located close to the NMJ that respond to nerve injury by engrafting to the post-synaptic compartment ^56^. Similarly, genetic ablation of MuSC has highlighted the important role of these cells in the maintenance and the regeneration of NMJ during aging and following sciatic nerve injury ^54,55^. Altogether these observations support a crosstalk between quiescent MuSC and motor nerve terminals. In line with this hypothesis, we showed that depletion of SMN-deficient MuSC induces a rapid remodeling of the NMJ evidenced by an increased proportion of poly-innervated NMJ, a hallmark of denervation/reinnervation occurring upon nerve injury, neurological pathologies and aging ^74–76^. At longer term, we found that SMN-deficient MuSC depletion leads to a non-cell autonomous loss of MN at all levels of the spinal cord. Intriguingly, in both cKO mouse models we used, although MuSC depletion was more pronounced in HMZ muscles than in HTZ muscles, the reduction in MN number was equivalent. Interestingly, single cell analysis of MN from SMA and control patients in culture revealed that SMN levels are highly variable and that MN expressing high levels of SMN display a better survival ^77^. According to available snRNAseq data, such heterogeneity in SMN level of expression also exists in the MuSC pool. We thus hypothesize that NMJ-associated MuSC may have higher SMN requirements and be preferentially targeted in HTZ muscles. Further investigation will be needed to clarify the biological significance of this heterogeneous expression of SMN and whether MuSC have differential SMN requirements. Nevertheless, since selective loss of αMN associated with sparing of γMN has also been reported in severe SMA mice ^61^, our data suggest that MuSC-targeted SMN deficit could reproduce some features of long-term SMA pathology.

NMJ are traditionally defined as a tripartite involving the muscle fiber, the motor neuron axon terminal and a terminal Schwann cell. Our findings show that the landscape might be more complex and that quiescent MuSC are also involved in NMJ maintenance. We hypothesize that in our model, MN loss might be either directly driven by the disappearance of trophic signals delivered by NMJ-associated MuSC or indirectly via muscle fiber defects consecutive to MuSC depletion. Our lab has recently provided a transcriptomic analysis of quiescent MuSC ^78^. From this study we identified several potential candidates that could elicit a paracrine effect on the motor neuron axon terminal. Among these secreted factors, Glial-derived neurotrophic factor (GDNF) might be of interest given its neuroprotective effects ^79,80^.

From a therapeutic point of view, our study demonstrates that MuSC pool attrition could contribute to the progression of SMA by exacerbating MN loss. Accordingly, in case of insufficient targeting of MuSC by current therapies, quiescent MuSC pool could be disrupted and, in turn, mediate MN loss at longer term. AAV-mediated *SMN1* replacement therapy is administered at a very early stage in children with SMA type I (6 to 9 months), a period during which postnatal myogenesis is active and myogenic progenitors are proliferating. Since the AAV vector transgene remains in episomal form in transduced cells, proliferating myogenic cells are likely to rapidly lose it and become deficient in SMN. Similarly, for patients treated at later stages, the pool of quiescent MuSC may already be disrupted, which in the long term could compromise the integrity of the neuromuscular system. Lastly, regarding patients treated by intrathecal delivery of nusinersen or oral administration of risdiplam the question arises as to whether MuSC are effectively targeted by these delivery routes. Therefore, our data highlight the importance of targeting MuSC effectively and early in the development of the pathology to preserve the integrity of the neuromuscular system over the long term and could therefore have major implications for the development of next generation therapeutic strategies.

From a broader perspective, our study highlights a strong interdependence between quiescent MuSC and MN reservoirs. Although the molecular mechanisms underlying this link remain to be identified, we anticipate that our findings could also be extended to the understanding of other neuromuscular myopathies affecting MuSC pool integrity^81^ and aging ^82^.

## STUDY APPROVAL

Animal handling and experimentation were performed according to French and European Community guidelines (Project agreement numbers: APAFIS#10477- 2017062616164326 and 30967-2021040615286805).

## AUTHOR CONTRIBUTIONS

JMec, JMig and ND conducted the experiments, acquired and analyzed the data. ND designed the research study. TO performed bioinformatic analysis. SA participated in animal experimentation and handling of mouse colonies. Jmig, MG, JB, SB and JMes analyzed the NMJ. ND, FR, MB, MGB and HR provided the fundings. Jmec and ND wrote the manuscript.

## ACKNOLEDGEMENTS

We thank: Matthew Borok for proofreading the manuscript, the MyoBank facility from the Institute of Myology for providing SMA patient muscle biopsies, C. Blanc and B. Hoareau from the Flow Cytometry Core CyPS (Sorbonne University, Pitié-Salpétrière Hospital, Paris, France). This work was supported by SMA Europe, AFM-Téléthon (Translamuscle Pole) and ANR.

## CONFLICT of INTEREST

The authors declare no conflict of interest.

## MATERIAL and METHODS

### Human muscles biopsies

Human muscle biopsies were provided by the AIM-MyoBank (Association Institut de Myologie, Pitié Salpétrière, Paris) and collected during surgical procedure in accordance with the French legislation on ethical rules. Written informed consent was obtained for all the biopsies used in this study.

### Mouse lines

Mouse lines used in this study were kindly provided by the corresponding laboratories or purchased from Jackson laboratory : *Pax7^CreERT2(Fan)^* (81), *Rosa26^mTmG^* ^57^, *_Pax7_CreERT2(Gaka)* 51_, *Rosa26*_*iDTR* 84 _and *Smn*_*F7/F7* _(50). *Pax7*_*CreERT2/+(Fan)* _mice were_ crossed with *Smn^F7/F7^* and *Rosa26^mTmG^* mice to generate the *Pax7^CreERT2/+^;Smn* floxed and the *Pax7^CreERT2/+^;R26^mTmG/+^* mouse lines. *Pax7^CreERT2(Gaka)^* mice were crossed with *Smn^F7/F7^* and *Rosa26^iDTR^*mice to obtain *Pax7^CreERT2/CreERT2^;Smn* floxed (referred as *_Pax7_CreERT2(Gaka)_;Smn_fl/fl*_) and *Pax7*_*CreERT2/CreERT2_;R26_iDTR/iDTR* _(referred as_ *Pax7^CreERT2(Gaka)^;R26^iDTR/iDTR^*) mice. All mice used in this study were 2-4 months old adult males. Nuclear translocation of the Cre recombinase was induced by intraperitoneal injections of Tamoxifen (Sigma-Aldrich) diluted in corn oil, for 4 consecutive days at the dose of 3mg/20g of mice body weight as previously described. ^85^

### Intramuscular delivery of scAAV9 vectors

Pseudotyped AAV9 vectors encoding *GFP* or *SMN1* transgenes under the control of PGK promoter were prepared by helper virus-free three-plasmid transfection of HEK293T cells, as previously described ^14^. 10^11^ vg of scAA9-GFP or scAAV9-SMN vectors were distributed intramuscularly in hindlimb muscles (Tibialis Anterior, Quadriceps and Gastrocnemius) from one-day old wildtype mice. Injected muscles were harvested and enzymatically digested or snap frozen, 7 days later for analysis.

### List of antibodies used for immunofluorescence

The following primary antibodies were used: mouse IgG1 anti-PAX7 (Developmental Studies Hybridoma Bank [DSHB]), rabbit anti-MYOD M-318 (Santa Cruz), mouse IgG1 anti-MYOG 5FD (Santa Cruz), rabbit anti-KI67 D3B5 (Cell Signaling), rabbit anti-Laminin (Sigma), mouse IgG2b anti-MyHC I BA-D5 (DSHB), mouse IgG1 anti-MyHC-IIa SC-71 (DSHB), mouse IgM anti-MyHC-IIb BF-F3 (DSHB), mouse IgM anti-MyHC IIx 6H1 (DSHB), rabbit anti-GFP (Abcam), goat anti-ChAT (Millipore), mouse IgG1 anti-NeuN (Millipore), mouse IgG1 anti-SMN (BD Biosciences), rabbit anti-SMN (Santa Cruz), chicken anti-Neurofilament (Millipore).

The following secondary antibodies were used: goat anti-mouse IgG1 biotinylated, Streptavidin-Cy3, goat anti-mouse IgG1-Cy3, goat anti-mouse IgG1-AF488, goat anti-rabbit-AF488, goat anti-chicken-AF488, goat anti-rabbit-Cy5 (all from Jackson Immunoresearch), donkey anti goat-AF488, goat anti mouse IgG2b-AF350 (Invitrogen), goat anti mouse IgM-AF647 (from Invitrogen).

### Analysis of MuSC apoptosis

Hindlimb muscles were harvested and dissociated with Collagenase/Dispase as previously described ^86^. Mononucleated cells from digested muscles were incubated for 45 min on ice with the following coupled antibodies: rat anti-mouse CD45-PECy7, TER-119-PECy7, CD31-PE-Cy7, CD34-bV421, SCA-1-FITC (BD Biosciences) and mouse anti-mouse ITGA7-AF700 (R&D Systems). Cells were washed twice with HBSS and then stained with coupled Annexin V from PE Annexin V apoptosis detection kit (BD Pharmingen), following manufacturer instructions. Analyses were performed using a BD FACS Canto^TM^ Flow Cytometer (BD Biosciences).

### Immunostainings on muscle sections

Muscles were dissected, embedded in OCT and snap frozen in liquid nitrogen-cooled isopentane. Muscle sections (10µm) were fixed with paraformaldehyde (4% PFA) for 20 min and permeabilized with cold methanol (-20°C) for 6 minutes. Antigen retrieval was performed by immersing the sections in 2 successive baths of hot citric acid (95°C) for 5 min followed by a resting phase of 5 min. Sections were then blocked for 2h with Bovine Serum Albumin (5% BSA, Jackson Immunoresearch) and then for 30 min with Goat anti-mouse immunoglobulin G (IgG) Fab fragment (Jackson Immunoresearch). Sections were incubated with primary antibodies overnight at 4°C or 2h at room temperature and then with appropriate coupled secondary antibodies for 45 min at room temperature. Slides were mounted with Fluoromount-G (Thermofisher).

### Muscle fiber phenotyping

For fiber phenotyping, serial muscle sections were incubated for 2h at room temperature with 5% BSA and for 30 min with goat anti-mouse immunoglobulin G (IgG) Fab fragment. Immunostaining was then proceeded as previously described ^87^. Briefly, muscle sections were incubated overnight at 4°C with primary antibodies anti-Laminin and anti-MyHC IIb (BF-F3) or MyHC IIx (6H1) and then with appropriate secondary antibodies for 45 min at room temperature. Next, sections were incubated with primary antibodies anti-MyHC I (BA-D5) and anti-MyHC IIa (SC71) for 2h at room temperature and corresponding secondary antibodies for 45 min. Slides were mounted with Fluoromount-G. Cross sectional area were determined using an Image J software plugin: MuscleJ ^88^.

### Immunostaining on spinal cord sections

Spines were dissected after intra cardiac perfusion of PBS and 4% paraformaldehyde (4% PFA) and incubated overnight at 4°C in 4% PFA. Spines were next immersed in 30% sucrose at 4°C for 24h prior to spinal cord recovery. Spinal cords were divided in three parts (cervical, thoracic and lumbar) embedded in OCT and snap frozen in liquid nitrogen-cooled isopentane. Cryosections (14µm) were fixed with 4% PFA for 30 min and permeabilized with 0.1% Triton X-100 (Sigma). Antigen retrieval was performed as described above for muscle sections. Next, spinal cord sections were blocked for 2h with a solution containing 5% BSA, 10% Donkey Serum and 0.1% Triton X-100 and for 30 min with goat anti-mouse immunoglobulin G (IgG) Fab fragment. Spinal cord sections were first incubated with primary antibody anti-ChAT overnight at 4°C and secondary antibody for 45 min. Sections were then incubated with primary antibody anti-NeuN for 2h at room temperature and the corresponding secondary antibody for 45 min. Slides were mounted with Fluoromount-G.

### Niels coloration of spinal cord sections

Spinal cord sections were stained for 15 min with Niels staining solution (0.1% Cresyl Violet acetate, 0.25% acetic acid). Sections were then washed twice with milliQ water and dehydrated by immersion in increasing concentrations of ethanol and three successive baths of xylene. Slides were mounted with Eukitt mounting medium (Sigma). Motor neurons were visualized by optic microscopy and counted according to their localization and shape.

### TA muscle clearing and NMJ immunostaining for 3D imaging

TA muscles were harvested and immediately fixed in 4% PFA for 2 hours at 4°C. Muscles were washed three times with PBS for 10 min at RT. After fixation, TA muscles were cut into 1 mm-thick slices longitudinally using a microtome. Samples were then permeabilized with 2% Triton-X 100 in PBS overnight at RT. Then, all the immunostaining steps were performed in dark conditions at 37°C under agitation at 10 rpm. Muscle slices were incubated with primary antibodies diluted in dilution buffer containing 1% goat serum, 0.2% Triton-X 100 and 0.2% sodium azide in PBS for 4-5 days at 37°C. Samples were extensively rinsed 3 times with PBS for at least 1 hour at RT, under agitation. Samples were then incubated with secondary antibodies for 2 days. For the last 2 hours, DAPI (1/500, Sigma) was added. After 4 washes of minimum 1 hour at RT, samples were processed for tissue clearing by incubation with RapiClear® 1.52 reagent (Nikon, France) for at least 12h. Imaging was performed on a Spinning disk microscope (Leica) using 25x water-immersion objective. Image processing was performed using Imaris 9 software. For NMJ innervation status determination, vacant motor endplates were considered as non-innervated NMJ. NMJ were considered “partially innervated” when the overlap between the nerve terminal and the motor endplate was ≤ 50%, and “fully innervated” when the overlap was > 50%. NMJ were defined as poly-innervated when innervated by at least 2 distinct terminal axons. Between 272 to 508 NMJ were analyzed per genotype (minimum 75 NMJ per mouse).

### Statistics

Results are presented as the mean±sem. All the statistical analyses were performed using GraphPad Prism software. Statistical significance was calculated by Student’s unpaired *t*-tests for single comparison, or by one-way and two-way ANOVA followed by post hoc tests specified in figure legends. *p*<0.05 was considered significant with *\*p<0.05*, \*\**p<0.01*, *\*\*\*p<0.001* and *\*\*\*\*p<0.0001*. Statistical analysis of mouse survival was determined using a log-rank test (Mantel-Cox).

## Notes

### Competing Interest Statement

The authors have declared no competing interest.

## REFERENCES

1. Kolb, S.J., and Kissel, J.T. (2015). Spinal Muscular Atrophy. Neurol. Clin. 33, 831–846. 10.1016/j.ncl.2015.07.004.

2. Lefebvre, S., Bürglen, L., Reboullet, S., Clermont, O., Burlet, P., Viollet, L., Benichou, B., Cruaud, C., Millasseau, P., Zeviani, M., et al. (1995). Identification and characterization of a spinal muscular atrophy-determining gene. Cell 80, 155–165. 10.1016/0092-8674(95)90460-3.

3. Monani, U.R. (1999). A single nucleotide difference that alters splicing patterns distinguishes the SMA gene SMN1 from the copy gene SMN2. Hum. Mol. Genet. 8, 1177–1183. 10.1093/hmg/8.7.1177.

4. Lorson, C.L., Hahnen, E., Androphy, E.J., and Wirth, B. (1999). A single nucleotide in the SMN gene regulates splicing and is responsible for spinal muscular atrophy. Proc. Natl. Acad. Sci. 96, 6307– 6311. 10.1073/pnas.96.11.6307.

5. Vitte, J., Fassier, C., Tiziano, F.D., Dalard, C., Soave, S., Roblot, N., Brahe, C., Saugier-Veber, P., Bonnefont, J.P., and Melki, J. (2007). Refined Characterization of the Expression and Stability of the SMN Gene Products. Am. J. Pathol. 171, 1269–1280. 10.2353/ajpath.2007.070399.

6. Campbell, L., Potter, A., Ignatius, J., Dubowitz, V., and Davies, K. (1997). Genomic Variation and Gene Conversion in Spinal Muscular Atrophy: Implications for Disease Process and Clinical Phenotype. Am. J. Hum. Genet. 61, 40–50. 10.1086/513886.

7. Calucho, M., Bernal, S., Alías, L., March, F., Venceslá, A., Rodríguez-Álvarez, F.J., Aller, E., Fernández, R.M., Borrego, S., Millán, J.M., et al. (2018). Correlation between SMA type and SMN2 copy number revisited: An analysis of 625 unrelated Spanish patients and a compilation of 2834 reported cases. Neuromuscul. Disord. 28, 208–215. 10.1016/j.nmd.2018.01.003.

8. Rao, V.K., Kapp, D., and Schroth, M. (2018). Gene Therapy for Spinal Muscular Atrophy: An Emerging Treatment Option for a Devastating Disease. J. Manag. Care Spec. Pharm. 24, S3–S16. 10.18553/jmcp.2018.24.12-a.s3.

9. Passini, M.A., Bu, J., Richards, A.M., Kinnecom, C., Sardi, S.P., Stanek, L.M., Hua, Y., Rigo, F., Matson, J., Hung, G., et al. (2011). Antisense oligonucleotides delivered to the mouse CNS ameliorate symptoms of severe spinal muscular atrophy. Sci. Transl. Med. 3, 72ra18. 10.1126/scitranslmed.3001777.

10. Finkel, R.S., Mercuri, E., Darras, B.T., Connolly, A.M., Kuntz, N.L., Kirschner, J., Chiriboga, C.A., Saito, K., Servais, L., Tizzano, E., et al. (2017). Nusinersen versus Sham Control in Infantile-Onset Spinal Muscular Atrophy. N. Engl. J. Med. 377, 1723–1732. 10.1056/NEJMoa1702752.

11. Ratni, H., Ebeling, M., Baird, J., Bendels, S., Bylund, J., Chen, K.S., Denk, N., Feng, Z., Green, L., Guerard, M., et al. (2018). Discovery of Risdiplam, a Selective Survival of Motor Neuron-2 ( *SMN2*) Gene Splicing Modifier for the Treatment of Spinal Muscular Atrophy (SMA). J. Med. Chem. 61, 6501–6517. 10.1021/acs.jmedchem.8b00741.

12. Darras, B.T., Masson, R., Mazurkiewicz-Bełdzińska, M., Rose, K., Xiong, H., Zanoteli, E., Baranello, G., Bruno, C., Vlodavets, D., Wang, Y., et al. (2021). Risdiplam-Treated Infants with Type 1 Spinal Muscular Atrophy versus Historical Controls. N. Engl. J. Med. 385, 427–435. 10.1056/NEJMoa2102047.

13. Valori, C.F., Ning, K., Wyles, M., Mead, R.J., Grierson, A.J., Shaw, P.J., and Azzouz, M. (2010). Systemic Delivery of scAAV9 Expressing SMN Prolongs Survival in a Model of Spinal Muscular Atrophy. Sci. Transl. Med. 2, 35ra42–35ra42. 10.1126/scitranslmed.3000830.

14. 14. Dominguez, E., Marais, T., Chatauret, N., Benkhelifa-Ziyyat, S., Duque, S., Ravassard, P., Carcenac, R., Astord, S., de Moura, A.P., Voit, T., et al. (2011). Intravenous scAAV9 delivery of a codon-optimized SMN1 sequence rescues SMA mice. Hum. Mol. Genet. 20, 681–693. 10.1093/hmg/ddq514.

15. Mendell, J.R., Al-Zaidy, S., Shell, R., Arnold, W.D., Rodino-Klapac, L.R., Prior, T.W., Lowes, L., Alfano, L., Berry, K., Church, K., et al. (2017). Single-Dose Gene-Replacement Therapy for Spinal Muscular Atrophy. N. Engl. J. Med. 377, 1713–1722. 10.1056/NEJMoa1706198.

16. Chaytow, H., Faller, K.M.E., Huang, Y.-T., and Gillingwater, T.H. (2021). Spinal muscular atrophy: From approved therapies to future therapeutic targets for personalized medicine. Cell Rep. Med. 2, 100346. 10.1016/j.xcrm.2021.100346.

17. Le, T.T., McGovern, V.L., Alwine, I.E., Wang, X., Massoni-Laporte, A., Rich, M.M., and Burghes, A.H.M. (2011). Temporal requirement for high SMN expression in SMA mice. Hum. Mol. Genet. 20, 3578–3591. 10.1093/hmg/ddr275.

18. Ramos, D.M., d’Ydewalle, C., Gabbeta, V., Dakka, A., Klein, S.K., Norris, D.A., Matson, J., Taylor, S.J., Zaworski, P.G., Prior, T.W., et al. (2019). Age-dependent SMN expression in disease-relevant tissue and implications for SMA treatment. J. Clin. Invest. 129, 4817–4831. 10.1172/JCI124120.

19. Mendell, J.R., Al-Zaidy, S.A., Lehman, K.J., McColly, M., Lowes, L.P., Alfano, L.N., Reash, N.F., Iammarino, M.A., Church, K.R., Kleyn, A., et al. (2021). Five-Year Extension Results of the Phase 1 START Trial of Onasemnogene Abeparvovec in Spinal Muscular Atrophy. JAMA Neurol. 78, 834. 10.1001/jamaneurol.2021.1272.

20. Gavrilina, T.O., McGovern, V.L., Workman, E., Crawford, T.O., Gogliotti, R.G., DiDonato, C.J., Monani, U.R., Morris, G.E., and Burghes, A.H.M. (2008). Neuronal SMN expression corrects spinal muscular atrophy in severe SMA mice while muscle-specific SMN expression has no phenotypic effect. Hum. Mol. Genet. 17, 1063–1075. 10.1093/hmg/ddm379.

21. 21. Martinez, T.L., Kong, L., Wang, X., Osborne, M.A., Crowder, M.E., Van Meerbeke, J.P., Xu, X., Davis, C., Wooley, J., Goldhamer, D.J., et al. (2012). Survival Motor Neuron Protein in Motor Neurons Determines Synaptic Integrity in Spinal Muscular Atrophy. J. Neurosci. 32, 8703–8715. 10.1523/JNEUROSCI.0204-12.2012.

22. Paez-Colasante, X., Seaberg, B., Martinez, T.L., Kong, L., Sumner, C.J., and Rimer, M. (2013). Improvement of Neuromuscular Synaptic Phenotypes without Enhanced Survival and Motor Function in Severe Spinal Muscular Atrophy Mice Selectively Rescued in Motor Neurons. PLoS ONE 8, e75866. 10.1371/journal.pone.0075866.

23. Hua, Y., Liu, Y.H., Sahashi, K., Rigo, F., Bennett, C.F., and Krainer, A.R. (2015). Motor neuron cell-nonautonomous rescue of spinal muscular atrophy phenotypes in mild and severe transgenic mouse models. Genes Dev. 29, 288–297. 10.1101/gad.256644.114.

24. Besse, A., Astord, S., Marais, T., Roda, M., Giroux, B., Lejeune, F.-X., Relaix, F., Smeriglio, P., Barkats, M., and Biferi, M.G. (2020). AAV9-Mediated Expression of SMN Restricted to Neurons Does Not Rescue the Spinal Muscular Atrophy Phenotype in Mice. Mol. Ther. J. Am. Soc. Gene Ther. 28, 1887–1901. 10.1016/j.ymthe.2020.05.011.

25. 25. Van Alstyne, M., Simon, C.M., Sardi, S.P., Shihabuddin, L.S., Mentis, G.Z., and Pellizzoni, L. (2018). Dysregulation of Mdm2 and Mdm4 alternative splicing underlies motor neuron death in spinal muscular atrophy. Genes Dev. 32, 1045–1059. 10.1101/gad.316059.118.

26. Hua, Y., Sahashi, K., Rigo, F., Hung, G., Horev, G., Bennett, C.F., and Krainer, A.R. (2011). Peripheral SMN restoration is essential for long-term rescue of a severe spinal muscular atrophy mouse model. Nature 478, 123–126. 10.1038/nature10485.

27. Hamilton, G., and Gillingwater, T.H. (2013). Spinal muscular atrophy: going beyond the motor neuron. Trends Mol. Med. 19, 40–50. 10.1016/j.molmed.2012.11.002.

28. 28. Nicole, S., Desforges, B., Millet, G., Lesbordes, J., Cifuentes-Diaz, C., Vertes, D., Cao, M.L., De Backer, F., Languille, L., Roblot, N., et al. (2003). Intact satellite cells lead to remarkable protection against Smn gene defect in differentiated skeletal muscle. J. Cell Biol. 161, 571–582. 10.1083/jcb.200210117.

29. Kim, J.-K., Jha, N.N., Feng, Z., Faleiro, M.R., Chiriboga, C.A., Wei-Lapierre, L., Dirksen, R.T., Ko, C.-P., and Monani, U.R. (2020). Muscle-specific SMN reduction reveals motor neuron–independent disease in spinal muscular atrophy models. J. Clin. Invest. 130, 1271–1287. 10.1172/JCI131989.

30. Kim, J.-H., Kang, J.-S., Yoo, K., Jeong, J., Park, I., Park, J.H., Rhee, J., Jeon, S., Jo, Y.-W., Hann, S.-H., et al. (2022). Bap1/SMN axis in Dpp4+ skeletal muscle mesenchymal cells regulates the neuromuscular system. JCI Insight 7, e158380. 10.1172/jci.insight.158380.

31. Hayhurst, M., Wagner, A.K., Cerletti, M., Wagers, A.J., and Rubin, L.L. (2012). A cell-autonomous defect in skeletal muscle satellite cells expressing low levels of survival of motor neuron protein. Dev. Biol. 368, 323–334. 10.1016/j.ydbio.2012.05.037.

32. 32. Liu, H., Chehade, L., Deguise, M.-O., De Repentigny, Y., and Kothary, R. (2024). SMN depletion impairs skeletal muscle formation and maturation in a mouse model of SMA. Hum. Mol. Genet.,ddae162. 10.1093/hmg/ddae162.

33. Rajendra, T.K., Gonsalvez, G.B., Walker, M.P., Shpargel, K.B., Salz, H.K., and Matera, A.G. (2007). A Drosophila melanogaster model of spinal muscular atrophy reveals a function for SMN in striated muscle. J. Cell Biol. 176, 831–841. 10.1083/jcb.200610053.

34. Walker, M.P., Rajendra, T.K., Saieva, L., Fuentes, J.L., Pellizzoni, L., and Matera, A.G. (2008). SMN complex localizes to the sarcomeric Z-disc and is a proteolytic target of calpain. Hum. Mol. Genet. 17, 3399–3410. 10.1093/hmg/ddn234.

35. Berciano, M.T., Castillo-Iglesias, M.S., Val-Bernal, J.F., Lafarga, V., Rodriguez-Rey, J.C., Lafarga, M., and Tapia, O. (2020). Mislocalization of SMN from the I-band and M-band in human skeletal myofibers in spinal muscular atrophy associates with primary structural alterations of the sarcomere. Cell Tissue Res. 381, 461–478. 10.1007/s00441-020-03236-3.

36. Cifuentes-Diaz, C., Frugier, T., Tiziano, F.D., Lacène, E., Roblot, N., Joshi, V., Moreau, M.H., and Melki, J. (2001). Deletion of murine SMN exon 7 directed to skeletal muscle leads to severe muscular dystrophy. J. Cell Biol. 152, 1107–1114. 10.1083/jcb.152.5.1107.

37. 37. van Haelst, U. (1970). An electron microscopic study of muscle in Werdnig-Hoffmann’s disease. Virchows Arch. A Pathol. Pathol. Anat. 351, 291–305. 10.1007/BF00547202.

38. Saito, Y. (1985). Muscle fibre type differentiation and satellite cell population in Werdnig-Hoffmann disease. J. Neurol. Sci. 68, 75–87. 10.1016/0022-510x(85)90051-6.

39. Martínez-Hernández, R., Soler-Botija, C., Also, E., Alias, L., Caselles, L., Gich, I., Bernal, S., and Tizzano, E.F. (2009). The developmental pattern of myotubes in spinal muscular atrophy indicates prenatal delay of muscle maturation. J. Neuropathol. Exp. Neurol. 68, 474–481. 10.1097/NEN.0b013e3181a10ea1.

40. Martínez-Hernández, R., Bernal, S., Alias, L., and Tizzano, E.F. (2014). Abnormalities in Early Markers of Muscle Involvement Support a Delay in Myogenesis in Spinal Muscular Atrophy. J. Neuropathol. Exp. Neurol. 73, 559–567. 10.1097/NEN.0000000000000078.

41. Guettier-Sigrist, S., Coupin, G., Braun, S., Rogovitz, D., Courdier, I., Warter, J.M., and Poindron, P. (2001). On the possible role of muscle in the pathogenesis of spinal muscular atrophy. Fundam. Clin. Pharmacol. 15, 31–40. 10.1046/j.1472-8206.2001.00006.x.

42. Shafey, D., Côté, P.D., and Kothary, R. (2005). Hypomorphic Smn knockdown C2C12 myoblasts reveal intrinsic defects in myoblast fusion and myotube morphology. Exp. Cell Res. 311, 49–61. 10.1016/j.yexcr.2005.08.019.

43. Bricceno, K.V., Martinez, T., Leikina, E., Duguez, S., Partridge, T.A., Chernomordik, L.V., Fischbeck, K.H., Sumner, C.J., and Burnett, B.G. (2014). Survival motor neuron protein deficiency impairs myotube formation by altering myogenic gene expression and focal adhesion dynamics. Hum. Mol. Genet. 23, 4745–4757. 10.1093/hmg/ddu189.

44. 44. Boyer, J.G., Deguise, M.-O., Murray, L.M., Yazdani, A., De Repentigny, Y., Boudreau-Larivière, C., and Kothary, R. (2014). Myogenic program dysregulation is contributory to disease pathogenesis in spinal muscular atrophy. Hum. Mol. Genet. 23, 4249–4259. 10.1093/hmg/ddu142.

45. Grandi, F.C., Astord, S., Gidaja, E., Pezet, S., Mazzucchi, S., Chapart, M., Vasseur, S., Mamchaoui, K., and Smeriglio, P. (2024). SMA Type II Skeletal Muscle Treated with Nusinersen shows SMN Restoration but Mitochondrial Deficiency. Preprint, 10.1101/2024.02.29.582680.

46. Zammit, P.S. (2006). Pax7 and myogenic progression in skeletal muscle satellite cells. J. Cell Sci., 1824–1832. 10.1242/jcs.02908.

47. Chaytow, H., Huang, Y.-T., Gillingwater, T.H., and Faller, K.M.E. (2018). The role of survival motor neuron protein (SMN) in protein homeostasis. Cell. Mol. Life Sci. 75, 3877–3894. 10.1007/s00018-018-2849-1.

48. Dachs, E., Hereu, M., Piedrafita, L., Casanovas, A., Calderó, J., and Esquerda, J.E. (2011). Defective Neuromuscular Junction Organization and Postnatal Myogenesis in Mice With Severe Spinal Muscular Atrophy. J. Neuropathol. Exp. Neurol. 70, 444–461. 10.1097/NEN.0b013e31821cbd8b.

49. Frugier, T., Tiziano, F.D., Cifuentes-Diaz, C., Miniou, P., Roblot, N., Dierich, A., Meur, M.L., and Melki, J. (2000). Nuclear targeting defect of SMN lacking the C-terminus in a mouse model of spinal muscular atrophy. Hum. Mol. Genet. 9, 849–858. 10.1093/hmg/9.5.849.

50. Lepper, C., Conway, S.J., and Fan, C.-M. (2009). Adult satellite cells and embryonic muscle progenitors have distinct genetic requirements. Nature 460, 627–631. 10.1038/nature08209.

51. Murphy, M.M., Lawson, J.A., Mathew, S.J., Hutcheson, D.A., and Kardon, G. (2011). Satellite cells, connective tissue fibroblasts and their interactions are crucial for muscle regeneration. Development 138, 3625–3637. 10.1242/dev.064162.

52. Mademtzoglou, D., Geara, P., Mourikis, P., and Relaix, F. (2023). Pax7 haploinsufficiency impairs muscle stem cell function in Cre-recombinase mice and underscores the importance of appropriate controls. Stem Cell Res. Ther. 14, 294. 10.1186/s13287-023-03506-1.

53. Dos Santos, M., Backer, S., Saintpierre, B., Izac, B., Andrieu, M., Letourneur, F., Relaix, F., Sotiropoulos, A., and Maire, P. (2020). Single-nucleus RNA-seq and FISH identify coordinated transcriptional activity in mammalian myofibers. Nat. Commun. 11, 5102. 10.1038/s41467-020-18789-8.

54. Liu, W., Wei-LaPierre, L., Klose, A., Dirksen, R.T., and Chakkalakal, J.V. (2015). Inducible depletion of adult skeletal muscle stem cells impairs the regeneration of neuromuscular junctions. eLife 4, e09221. 10.7554/eLife.09221.

55. Liu, W., Klose, A., Forman, S., Paris, N.D., Wei-LaPierre, L., Cortés-Lopéz, M., Tan, A., Flaherty, M., Miura, P., Dirksen, R.T., et al. (2017). Loss of adult skeletal muscle stem cells drives age-related neuromuscular junction degeneration. eLife 6, e26464. 10.7554/eLife.26464.

56. Larouche, J.A., Mohiuddin, M., Choi, J.J., Ulintz, P.J., Fraczek, P., Sabin, K., Pitchiaya, S., Kurpiers, S.J., Castor-Macias, J., Liu, W., et al. (2021). Murine muscle stem cell response to perturbations of the neuromuscular junction are attenuated with aging. eLife 10, e66749. 10.7554/eLife.66749.

57. Muzumdar, M.D., Tasic, B., Miyamichi, K., Li, L., and Luo, L. (2007). A global double-fluorescent Cre reporter mouse. genesis 45, 593–605. 10.1002/dvg.20335.

58. Kanning, K.C., Kaplan, A., and Henderson, C.E. (2010). Motor Neuron Diversity in Development and Disease. Annu. Rev. Neurosci. 33, 409–440. 10.1146/annurev.neuro.051508.135722.

59. Friese, A., Kaltschmidt, J.A., Ladle, D.R., Sigrist, M., Jessell, T.M., and Arber, S. (2009). Gamma and alpha motor neurons distinguished by expression of transcription factor Err3. Proc. Natl. Acad. Sci. 106, 13588–13593. 10.1073/pnas.0906809106.

60. Shneider, N.A., Brown, M.N., Smith, C.A., Pickel, J., and Alvarez, F.J. (2009). Gamma motor neurons express distinct genetic markers at birth and require muscle spindle-derived GDNF for postnatal survival. Neural Develop. 4, 42. 10.1186/1749-8104-4-42.

61. Powis, R.A., and Gillingwater, T.H. (2016). Selective loss of alpha motor neurons with sparing of gamma motor neurons and spinal cord cholinergic neurons in a mouse model of spinal muscular atrophy. J. Anat. 228, 443–451. 10.1111/joa.12419.

62. Jha, N.N., Kim, J.-K., Her, Y.-R., and Monani, U.R. (2023). Muscle: an independent contributor to the neuromuscular spinal muscular atrophy disease phenotype. JCI Insight 8, e171878. 10.1172/jci.insight.171878.

63. Ikenaka, A., Kitagawa, Y., Yoshida, M., Lin, C.-Y., Niwa, A., Nakahata, T., and Saito, M.K. (2023). SMN promotes mitochondrial metabolic maturation during myogenesis by regulating the MYOD- miRNA axis. Life Sci. Alliance 6, e202201457. 10.26508/lsa.202201457.

64. Berciano, M.T., Puente-Bedia, A., Medina-Samamé, A., Rodríguez-Rey, J.C., Calderó, J., Lafarga, M., and Tapia, O. (2020). Nusinersen ameliorates motor function and prevents motoneuron Cajal body disassembly and abnormal poly(A) RNA distribution in a SMA mouse model. Sci. Rep. 10, 10738. 10.1038/s41598-020-67569-3.

65. Keefe, A.C., Lawson, J.A., Flygare, S.D., Fox, Z.D., Colasanto, M.P., Mathew, S.J., Yandell, M., and Kardon, G. (2015). Muscle stem cells contribute to myofibres in sedentary adult mice. Nat. Commun. 6, 7087. 10.1038/ncomms8087.

66. 66. Sambasivan, R., Yao, R., Kissenpfennig, A., Van Wittenberghe, L., Paldi, A., Gayraud-Morel, B., Guenou, H., Malissen, B., Tajbakhsh, S., and Galy, A. (2011). Pax7-expressing satellite cells are indispensable for adult skeletal muscle regeneration. Development 138, 3647–3656. 10.1242/dev.067587.

67. Ando, S., Tanaka, M., Chinen, N., Nakamura, S., Shimazawa, M., and Hara, H. (2020). SMN Protein Contributes to Skeletal Muscle Cell Maturation Via Caspase-3 and Akt Activation. Vivo Athens Greece 34, 3247–3254. 10.21873/invivo.12161.

68. Hellbach, N., Peterson, S., Haehnke, D., Shankar, A., LaBarge, S., Pivaroff, C., Saenger, S., Thomas, C., McCarthy, K., Ebeling, M., et al. (2018). Impaired myogenic development, differentiation and function in hESC-derived SMA myoblasts and myotubes. PLOS ONE 13, e0205589. 10.1371/journal.pone.0205589.

69. Luchetti, A., Ciafrè, S.A., Murdocca, M., Malgieri, A., Masotti, A., Sanchez, M., Farace, M.G., Novelli, G., and Sangiuolo, F. (2015). A Perturbed MicroRNA Expression Pattern Characterizes Embryonic Neural Stem Cells Derived from a Severe Mouse Model of Spinal Muscular Atrophy (SMA). Int. J. Mol. Sci. 16, 18312–18327. 10.3390/ijms160818312.

70. Murdocca, M., Ciafrè, S.A., Spitalieri, P., Talarico, R.V., Sanchez, M., Novelli, G., and Sangiuolo, F. (2016). SMA Human iPSC-Derived Motor Neurons Show Perturbed Differentiation and Reduced miR-335-5p Expression. Int. J. Mol. Sci. 17. 10.3390/ijms17081231.

71. Schrank, B., Götz, R., Gunnersen, J.M., Ure, J.M., Toyka, K.V., Smith, A.G., and Sendtner, M. (1997). Inactivation of the survival motor neuron gene, a candidate gene for human spinal muscular atrophy, leads to massive cell death in early mouse embryos. Proc. Natl. Acad. Sci. U. S. A. 94, 9920–9925. 10.1073/pnas.94.18.9920.

72. Anderton, R.S., Meloni, B.P., Mastaglia, F.L., and Boulos, S. (2013). Spinal muscular atrophy and the antiapoptotic role of survival of motor neuron (SMN) protein. Mol. Neurobiol. 47, 821–832. 10.1007/s12035-013-8399-5.

73. 73. Sansa, A., de la Fuente, S., Comella, J.X., Garcera, A., and Soler, R.M. (2021). Intracellular pathways involved in cell survival are deregulated in mouse and human spinal muscular atrophy motoneurons. Neurobiol. Dis., 105366. 10.1016/j.nbd.2021.105366.

74. Bermedo-García, F., Zelada, D., Martínez, E., Tabares, L., and Henríquez, J.P. (2022). Functional regeneration of the murine neuromuscular synapse relies on long-lasting morphological adaptations. BMC Biol. 20, 158. 10.1186/s12915-022-01358-4.

75. Sleigh, J.N., Grice, S.J., Burgess, R.W., Talbot, K., and Cader, M.Z. (2014). Neuromuscular junction maturation defects precede impaired lower motor neuron connectivity in Charcot-Marie-Tooth type 2D mice. Hum. Mol. Genet. 23, 2639–2650. 10.1093/hmg/ddt659.

76. Blasco, A., Gras, S., Mòdol-Caballero, G., Tarabal, O., Casanovas, A., Piedrafita, L., Barranco, A., Das, T., Pereira, S.L., Navarro, X., et al. (2020). Motoneuron deafferentation and gliosis occur in association with neuromuscular regressive changes during ageing in mice. J. Cachexia Sarcopenia Muscle 11, 1628–1660. 10.1002/jcsm.12599.

77. Rodriguez-Muela, N., Litterman, N.K., Norabuena, E.M., Mull, J.L., Galazo, M.J., Sun, C., Ng, S.- Y., Makhortova, N.R., White, A., Lynes, M.M., et al. (2017). Single-Cell Analysis of SMN Reveals Its Broader Role in Neuromuscular Disease. Cell Rep. 18, 1484–1498. 10.1016/j.celrep.2017.01.035.

78. 78. Machado, L., Esteves de Lima, J., Fabre, O., Proux, C., Legendre, R., Szegedi, A., Varet, H., Ingerslev, L.R., Barrès, R., Relaix, F., et al. (2017). In Situ Fixation Redefines Quiescence and Early Activation of Skeletal Muscle Stem Cells. Cell Rep. 21, 1982–1993. 10.1016/j.celrep.2017.10.080.

79. Oppenheim, R.W., Houenou, L.J., Johnson, J.E., Lin, L.F., Li, L., Lo, A.C., Newsome, A.L., Prevette, D.M., and Wang, S. (1995). Developing motor neurons rescued from programmed and axotomy-induced cell death by GDNF. Nature 373, 344–346. 10.1038/373344a0.

80. Angka, H.E., Geddes, A.J., and Kablar, B. (2008). Differential survival response of neurons to exogenous GDNF depends on the presence of skeletal muscle. Dev. Dyn. Off. Publ. Am. Assoc. Anat. 237, 3169–3178. 10.1002/dvdy.21727.

81. Ganassi, M., and Zammit, P.S. (2022). Involvement of muscle satellite cell dysfunction in neuromuscular disorders: Expanding the portfolio of satellite cell-opathies. Eur. J. Transl. Myol. 32. 10.4081/ejtm.2022.10064.

82. Chakkalakal, J.V., Jones, K.M., Basson, M.A., and Brack, A.S. (2012). The aged niche disrupts muscle stem cell quiescence. Nature 490, 355–360. 10.1038/nature11438.

83. Lepper, C., Conway, S.J., and Fan, C.-M. (2009). Adult satellite cells and embryonic muscle progenitors have distinct genetic requirements. Nature 460, 627–631. 10.1038/nature08209.

84. Buch, T., Heppner, F.L., Tertilt, C., Heinen, T.J.A.J., Kremer, M., Wunderlich, F.T., Jung, S., and Waisman, A. (2005). A Cre-inducible diphtheria toxin receptor mediates cell lineage ablation after toxin administration. Nat. Methods 2, 419–426. 10.1038/nmeth762.

85. Hayashi, S., and McMahon, A.P. (2002). Efficient recombination in diverse tissues by a tamoxifen-inducible form of Cre: a tool for temporally regulated gene activation/inactivation in the mouse. Dev. Biol. 244, 305–318. 10.1006/dbio.2002.0597.

86. Gattazzo, F., Laurent, B., Relaix, F., Rouard, H., and Didier, N. (2020). Distinct Phases of Postnatal Skeletal Muscle Growth Govern the Progressive Establishment of Muscle Stem Cell Quiescence. Stem Cell Rep. 15, 597–611. 10.1016/j.stemcr.2020.07.011.

87. Didier, N., Hourdé, C., Amthor, H., Marazzi, G., and Sassoon, D. (2012). Loss of a single allele for Ku80 leads to progenitor dysfunction and accelerated aging in skeletal muscle. EMBO Mol. Med. 4, 910–923. 10.1002/emmm.201101075.

88. Mayeuf-Louchart, A., Hardy, D., Thorel, Q., Roux, P., Gueniot, L., Briand, D., Mazeraud, A., Bouglé, A., Shorte, S.L., Staels, B., et al. (2018). MuscleJ: a high-content analysis method to study skeletal muscle with a new Fiji tool. Skelet. Muscle 8, 25. 10.1186/s13395-018-0171-0.

