## Supplemental information for "Targeted-SMN insufficiency in Skeletal Muscle Stem Cells mediates non-cell autonomous loss of motor neurons at long term"

**Mecca_Fig S1**

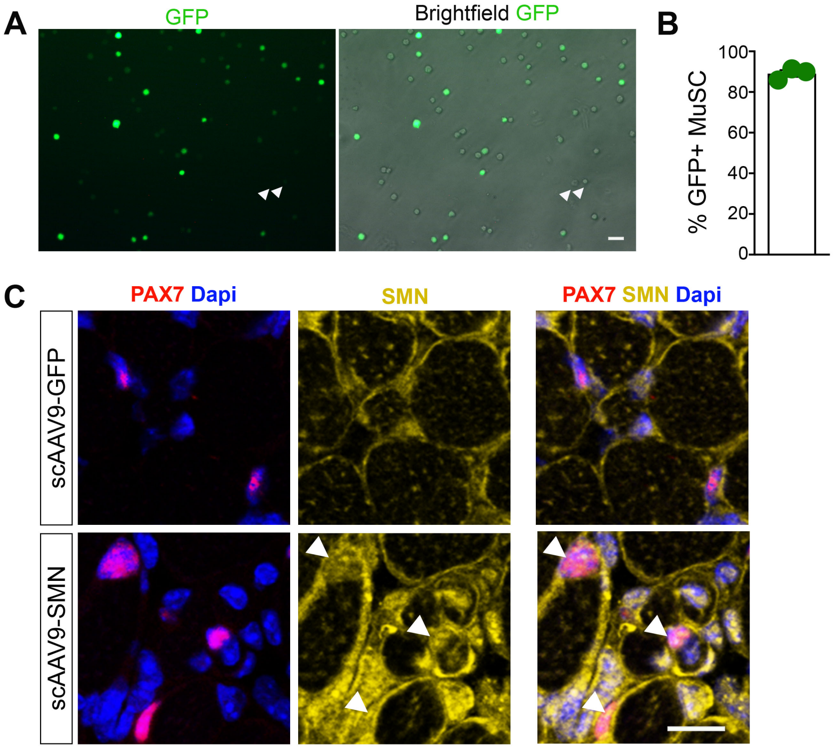
A
GFP
Brightfield GFP
B
% GFP+ MuSC
100
80
60
40
20
0
C
PAX7 Dapi
SMN
PAX7 SMN Dapi
scAAV9-GFP
scAAV9-SMN

**Mecca_Fig S2**

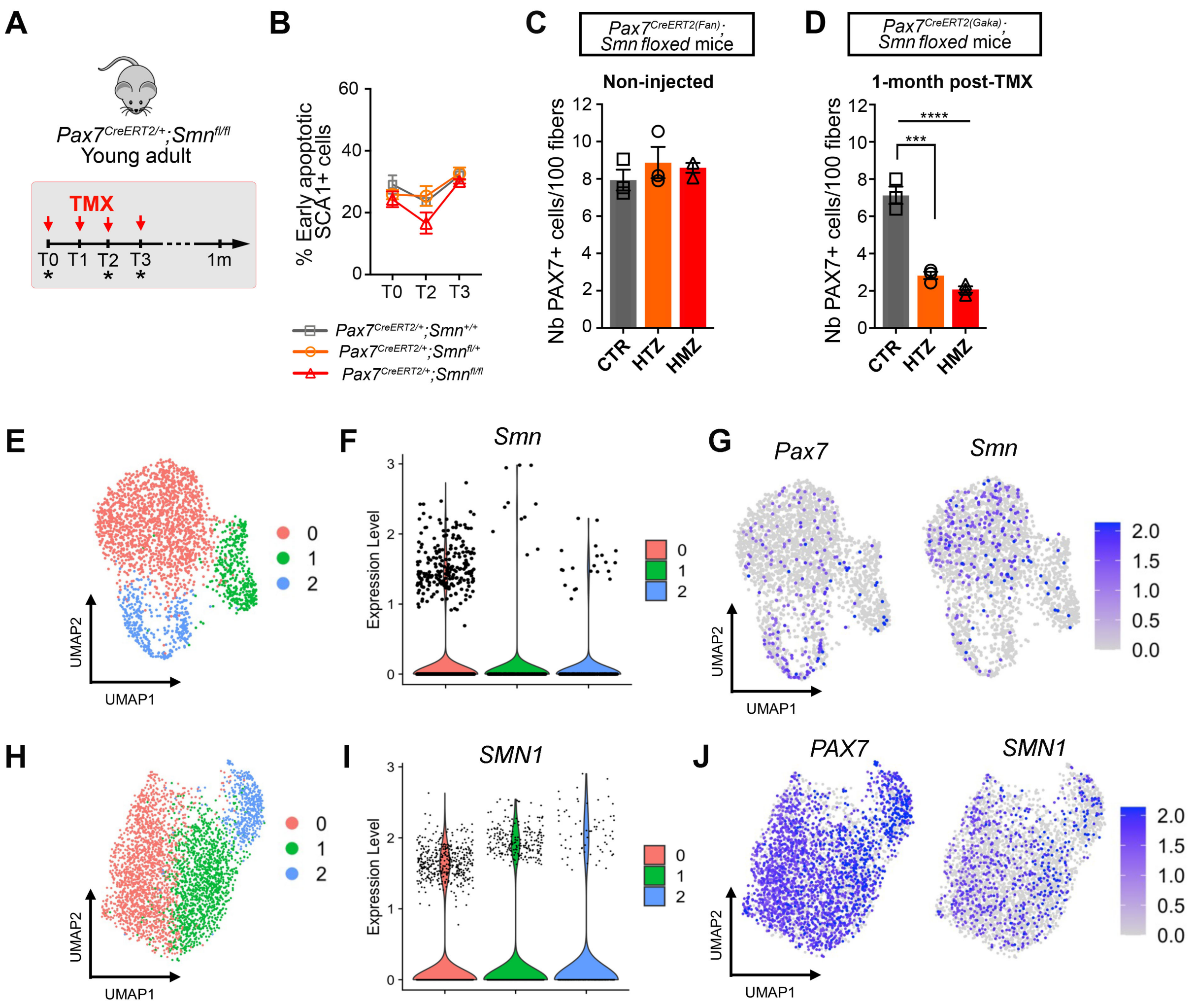

A
Pax7CreERT2/+;Smnfl/fl
Young adult
TMX
T0 T1 T2 T3 1m
B
% Early apoptotic SCA1+ cells
Pax7CreERT2/+;Smn+/+
Pax7CreERT2/+;Smnfl/+
Pax7CreERT2/+;Smnfl/fl
C
Pax7CreERT2(Fan);
Smn floxed mice
Non-injected
Nb PAX7+ cells/100 fibers
CTR
HTZ
HMZ
D
Pax7CreERT2(Gaka);
Smn floxed mice
1-month post-TMX
****
***
E
UMAP1
UMAP2
F
Smn
Expression Level
G
Pax7
Smn
H
I
SMN1
J
PAX7
SMN1

### Mecca_Fig S3

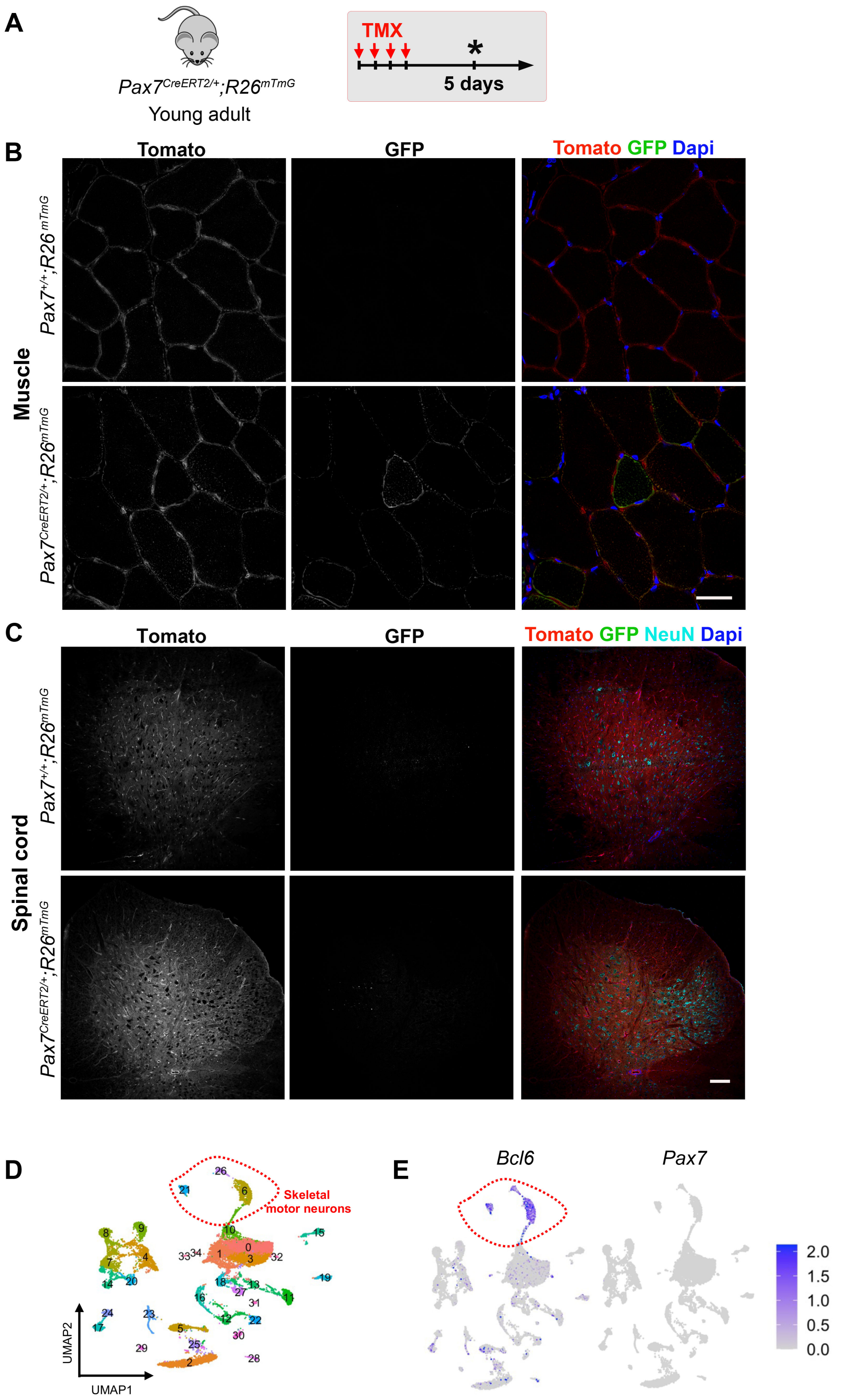
A
Pax7CreERT2/+;R26mTmG
Young adult
TMX
5 days
B
Tomato
GFP
Tomato GFP Dapi
Muscle
Pax7+/+;R26mTmG
Pax7CreERT2/+;R26mTmG
C
Tomato
GFP
Tomato GFP NeuN Dapi
Spinal cord
Pax7+/+;R26mTmG
Pax7CreERT2/+;R26mTmG
D
Skeletal motor neurons
UMAP1
UMAP2
E
Bcl6
Pax7
2.0
1.5
1.0
0.5
0.0

Mecca_Fig S4

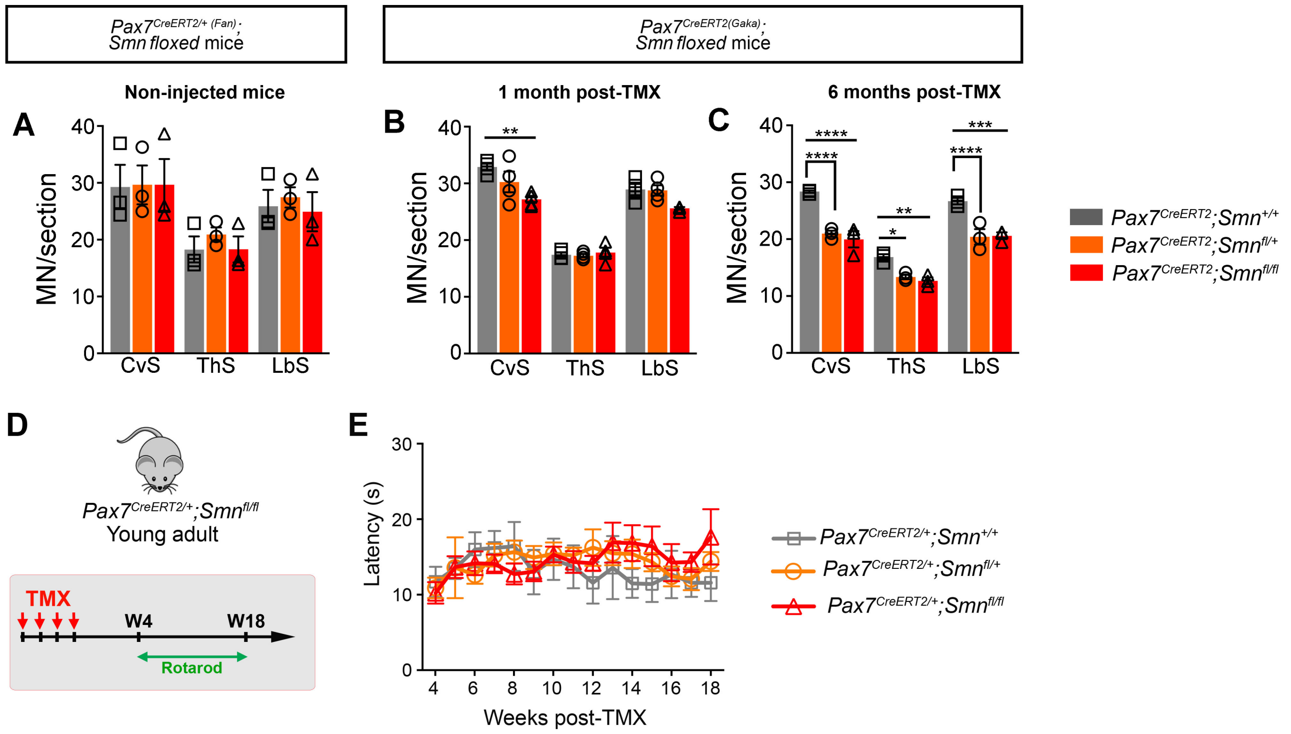

Pax7CreERT2/+ (Fan); Smn floxed mice
Pax7CreERT2(Gaka); Smn floxed mice
Non-injected mice
1 month post-TMX
6 months post-TMX
A
B
C
MN/section
CvS
ThS
LbS
Pax7CreERT2;Smn+/+
Pax7CreERT2;Smnfl/+
Pax7CreERT2;Smnfl/fl
D
Pax7CreERT2/+;Smnfl/fl
Young adult
TMX
W4
W18
Rotarod
E
Latency (s)
Weeks post-TMX
Pax7CreERT2/+;Smn+/+
Pax7CreERT2/+;Smnfl/+
Pax7CreERT2/+;Smnfl/fl

**Mecca_Fig S5**

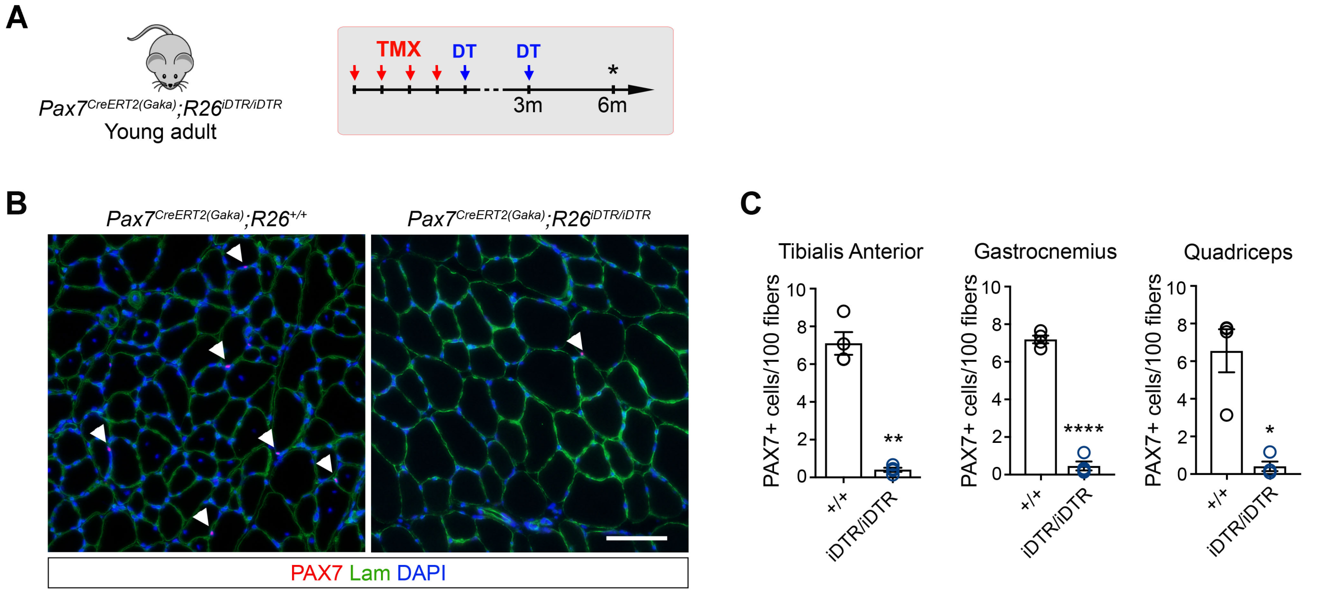
A
Pax7CreERT2(Gaka);R26iDTR/iDTR
Young adult
TMX
DT
DT
*
3m
6m
B
Pax7CreERT2(Gaka);R26+/+
Pax7CreERT2(Gaka);R26iDTR/iDTR
PAX7 Lam DAPI
C
Tibialis Anterior
Gastrocnemius
Quadriceps
PAX7+ cells/100 fibers
10
8
6
4
2
0
**
****
*
+/+
iDTR/iDTR

#### SUPPLEMENTAL FIGURE LEGENDS

**Figure S1 related to figure 2: scAAV9 vectors delivered intramuscularly mediate efficient transduction of MuSC in postnatal muscles.** $10^{11}$vg of scAA9-GFP vector were distributed intramuscularly in hindlimb muscles (Tibialis Anterior, Quadriceps and Gastrocnemius) of one day old WT mice. After 7 days, muscles were harvested, enzymatically digested and MuSC (CD45-Ter119-SCA1-CD34+ITGA7+) were purified by FACS. Purified MuSC were cytospun and GFP expression was assessed. (**A**) Images showing GFP expression in freshly isolated MuSC. Arrowheads point to GFP-MuSC. (**B**) Quantification of the proportion of GFP+ MuSC. (**C**) Representative images of TA muscle cross-sections, 7 days after injection with scAAV9-GFP or scAAV9-SMN vectors, immunostained for SMN and PAX7. Nuclei were stained with DAPI. Scale bars: 20 μm.

**Figure S2 related to figure 3: *Pax7 CreERT2* driven ablation of *Smn* specifically induces MuSC pool depletion in cKO mouse muscles.** (**A**) Chronograph of the experiment. (**B**) Percentage of early apoptotic SCA1+ cells in mononucleated cells from CTR, HTZ and HMZ mouse muscles determined by flow cytometry. (**C**) Number of PAX7+ MuSC per 100 fibers quantified on muscles sections from non-injected mice. (**D**) Number of PAX7+ MuSC per 100 fibers, quantified on muscle sections from *Pax7*$^{CreERT2(Gaka)}$*;Smn*$^{+/+}$ (CTR), *Pax7*$^{CreERT2(Gaka)}$*;Smn*$^{fl/+}$ (HTZ) and *Pax7*$^{CreERT2(Gaka)}$*;Smn*$^{fl/fl}$ (HMZ), 1-month post-TMX. Data are the mean±sem of 3 independent quantifications (n=3 mice/group). One-way ANOVA and Tukey's multiple comparisons test with ***$p<0.001$ and ****$p<0.0001$. (**E**) Uniform Manifold Approximation and Projection (UMAP) plot of different mouse MuSC subclusters based on single cell RNAseq data from [1]. (**F**) Violin plot of *Smn* expression in mouse MuSC subclusters. (**G**) Feature plot analysis of *Pax7* and *Smn* expression in mouse MuSC.

(**H**) UMAP plot of different human MuSC subclusters based on single cell RNAseq data from [2]. (**I**) Violin plot of *SMN1* gene expression in human MuSC subclusters. (**J**) Feature plot analysis of *PAX7* and *SMN1* expression in human MuSC.

**Figure S3 related to figures 5 and 6: Specificity of *Pax7 CreERT2* driven recombination in muscle tissue.** (**A**) Chronograph of the experiment. (**B, C**) Representative sections of TA muscles immunostained for GFP (**B**) and of spinal cords immunostained for GFP and NeuN (**C**), from adult $Pax7^{+/+};R26^{mTmG}$ and $Pax7^{CreERT2/+};R26^{mTmG}$ mice, 5 days after TMX injections. Scale bars: 100 μm. (**D**) UMAP plot of different MN subclusters previously identified in mouse spinal cord by single nuclei RNAseq from [3]. (**E**) Feature plot analysis of *Bcl6* and *Pax7* gene expression, showing the absence of expression of *Pax7* gene in αMN.

**Figure S4 related to figure 6: Depletion of SMN-deficient MuSC induces loss of MN in $Pax7^{CreERT2(Gaka)};Smn^{fl/+}$ and $Pax7^{CreERT2(Gaka)};Smn^{fl/fl}$ mice, at long term.** (**A**) Quantification of the number of MN/section on spinal cord sections from non-injected $Pax7^{CreERT2/+};Smn^{+/+}$, $Pax7^{CreERT2/+};Smn^{fl/+}$ and $Pax7^{CreERT2/+};Smn^{fl/fl}$ mice stained with Niels coloration. (**B, C**) Quantification of the number of MN/section on spinal cord sections from $Pax7^{CreERT2(Gaka)};Smn^{+/+}$, $Pax7^{CreERT2(Gaka)};Smn^{fl/+}$ and $Pax7^{CreERT2(Gaka)};Smn^{fl/fl}$ mice stained with Niels coloration, 1-month (**B**) and 6-months post-TMX (**C**). Data are the mean±sem of minimum 3 independent experiments (3-4 mice/group). Two-way ANOVA and Tukey's multiple comparisons test with *$p<0.05$, **$p<0.01$, ***$p<0.001$ and ****$p<0.0001$. (**D**) Experiment chronograph. Coordination of the mice was weekly evaluated by rotarod test with a progressive increase of the speed between 4 and 18 weeks post-TMX. (**E**) Line charts showing the mean latency to fall from rotarod in seconds (s). Values are the means±sem of 5-6 mice/group.

**Figure S5 related to figure 6: Efficacy of MuSC ablation in localized hindlimb muscles of *Pax7$^{CreERT2(Gaka)}$;R26$^{iDTR/iDTR}$* mice.** (**A**) Experiment chronograph. Diphteria Toxin (DT) was injected intramuscularly in Tibialis anterior, Gastrocnemius and Quadriceps muscles of adult *Pax7$^{CreERT2(Gaka)}$;R26$^{+/+}$* and *Pax7$^{CreERT2(Gaka)}$;R26$^{iDTR/iDTR}$* mice, the day after TMX injections and 3 months later. Muscles were analyzed 6 months post-TMX. (**B**) Representative images of Tibialis anterior muscle cross-sections immunostained for PAX7 and Laminin, 6 months post-TMX. (**C**) Quantification of the number of PAX7+ MuSC/100 fibers.

#### SUPPLEMENTAL REFERENCES


1. Dell'Orso, S., Juan, A.H., Ko, K.-D., Naz, F., Perovanovic, J., Gutierrez-Cruz, G., Feng, X., and Sartorelli, V. (2019). Single cell analysis of adult mouse skeletal muscle stem cells in homeostatic and regenerative conditions. Development *146*, dev174177. https://doi.org/10.1242/dev.174177.

2. Kedlian, V.R., Wang, Y., Liu, T., Chen, X., Bolt, L., Tudor, C., Shen, Z., Fasouli, E.S., Prigmore, E., Kleshchevnikov, V., et al. (2024). Human skeletal muscle aging atlas. Nat Aging *4*, 727–744. https://doi.org/10.1038/s43587-024-00613-3.

3. Blum, J.A., Klemm, S., Shadrach, J.L., Guttenplan, K.A., Nakayama, L., Kathiria, A., Hoang, P.T., Gautier, O., Kaltschmidt, J.A., Greenleaf, W.J., et al. (2021). Single-cell transcriptomic analysis of the adult mouse spinal cord reveals molecular diversity of autonomic and skeletal motor neurons. Nat Neurosci *24*, 572–583. https://doi.org/10.1038/s41593-020-00795-0.
